# Coiled-coil scaffolds for Targeted Protein Degradation

**DOI:** 10.64898/2025.12.22.695764

**Authors:** Boguslawa Korona, Bram Mylemans, Amanda M. Acevedo-Jake, Danny T. Huang, Thomas A. Edwards, Andrew J. Wilson, Derek N. Woolfson, Laura S. Itzhaki

**Author notes:** **Author contributions:** BK, BM, DH, TAE, AJW, DNW and LSI conceived the project and designed the experiments. BK generated all biological materials and performed all cell assays. BM designed the scaffolds, and BM and AMAJ designed the binders. BK and LSI wrote the paper. All authors have read and contributed to the preparation of the manuscript. LSI, DNW, AJW, DTH, and TAE raised the funding.

## Abstract

In targeted protein degradation (TPD) the cell’s quality control machinery is co-opted to degrade proteins of interest. Currently TPD is limited by the availability of small-molecule binders for targets and the many potential human E3 ubiquitin ligases. Consequently, other approaches are needed to tackle new targets and to exploit degradation pathways fully. The natural binding epitopes for E3s and other degradation machineries are often short linear peptide motifs, which offer routes to new biologics-based degraders. Here, targeting the anti-apoptotic BCL-x_L_ and initiation of apoptosis, we show that several protein-degradation pathways—ubiquitination, direct proteasome, autophagy-lysosome recruitment—can be leveraged by integrating target-binding and degradation motifs into single-polypeptide constructs. This exploits versatile and varied *de novo* coiled-coil scaffolds as adaptable molecular glues by presenting motifs multivalently and in diverse arrangements. The constructs are small (≈80 residues) and provide a plug-and-play platform to map and optimise target–degradation space using rational design. We call this <u>Co</u>iled C<u>o</u>ils for induced <u>Prox</u>imity (CoCoProx) and propose its use for targeted degradation and other induced-proximity reactions.

## Main

Targeted protein degradation (TPD), in which the cell’s quality control machinery is co-opted to drive the degradation of a disease-associated target, is a therapeutic strategy that has generated huge excitement in both academia and industry ^1 2 3^. TPD has been most successfully exemplified using small-molecule PROTACs (proteolysis targeting chimeras) composed of a target-binding ligand and a E3 ubiquitin ligase-binding ligand connected by a linker; there are now several PROTACs in clinical trials ^4, 5^. The TPD concept also encompasses molecular glues (ligands that stabilize the interaction between an E3 ubiquitin ligase (subsequently referred to as E3) and a neo substrate ^6, 7^, and it has been broadened to harness the autophagy-lysosome and endo-lysosome pathways, enabling larger macromolecules (such as protein aggregates) and membrane-associated and secreted proteins to be targeted ^8–11^.

The intense interest in TPD arises because: (i) target degradation has the potential to be longer lasting than target inhibition; (ii) degraders can be more potent than inhibitors given that one molecule can act on multiple targets; (iii) degraders should inhibit all functions of multi-functional targets, and, unlike conventional inhibitors, they do not require the ligand to block a functional site so have the potential to expand the scope of binding moieties to hard-to-drug targets; and (iv) as they mediate their effects through event-driven pharmacology, they offer potential selectivity and dose-limiting advantages because they rely on ternary complex formation^12, 13^. Nevertheless, degraders in the public domain are for targets that have already been drugged, leaving much of the proteome still to be accessed. Moreover, currently in TPD only a small fraction of the hundreds of human E3s have been harnessed, for example, Cereblon (CRBN) and von Hippel-Lindau (VHL) ^14–18^; and most do not have natural small-molecule ligands and/or are not readily amenable to ligand identification and optimization, as they tend to be multi-subunit, multi-domain protein complexes. Consequently, new approaches are required to tackle important therapeutic targets and harness degradation pathways. Biologics-based degraders, using peptide ligands, fill this lacuna. Given that many natural binding epitopes for E3s (referred to as “degrons”) are short linear interacting motifs (SLiMs) ^19–21^, biologic-based degraders should enable access to many more E3s than is possible with small molecules as well as to other degradation machineries, many of which also recognise SLiM-like epitopes on their substrates. These include degron-like sequences that label proteins for degradation by the autophagy-lysosome pathway, such as LIRs (LC3-interaction regions, also referred to as AIMs (Atg8-interacting motifs)) that are recognized by the LC3/GABARAP family of selective autophagy receptors ^22^. Lastly, peptide motifs can be exploited to direct targets to the proteasome potentially bypassing the need for ubiquitination by co-opting mechanisms such as “shuttle” proteins and natural “destabilization domains” ^23–25^.

Here we show how several of the cell’s protein degradation pathways can be accessed using a biologics approach (**Fig. 1a**), and we demonstrate how target-binding and degradation ligands can be integrated within single polypeptide chains using bifunctional *de novo* coiled-coil modules ^26^. By exploiting *de novo* designed coiled coils in varied formats (parallel or antiparallel dimers, trimers, and tetramers), we present the ligands in a multivalent manner to increase potency and enable diverse geometries to be tested. Previously, we have shown that such coiled coils can be engineered with new binding functions, making them ideal architectures to explore the concept of multi-valent and multi-functional proximity-inducing scaffolds ^27–31^.

**Fig. 1.**
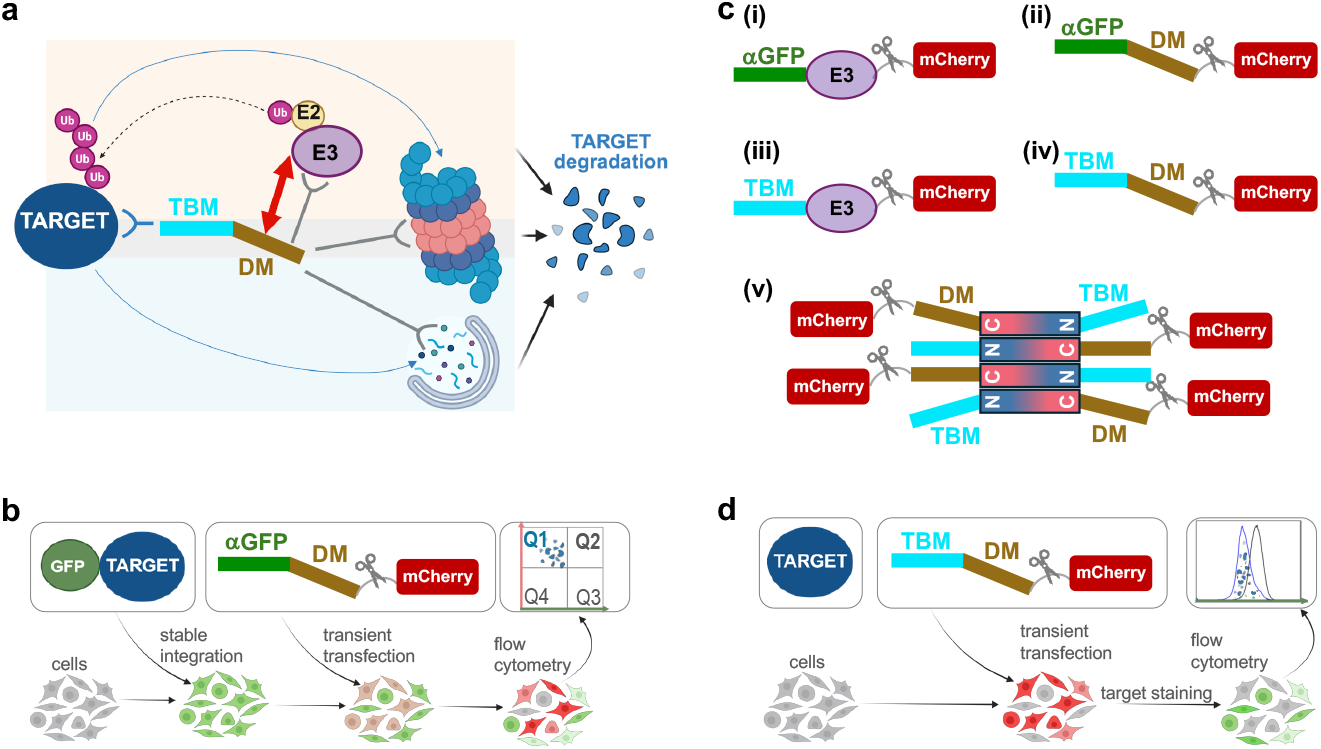
Schematics showing degrader design and assays. See Table S1 for amino-acid sequences. **(a)** Design of degraders comprising a target-binding motif (TBM) fused via a short flexible linker, composed of GS stretches, to a degradation motif (DM). DMs were designed for recruiting E3s, for direct proteasome recruitment, or for recruitment of the autophagy-lysosome pathway. **(b)** Schematic of flow cytometry-based degradation assay using a cell line stably expressing GFP-target fusion protein and transient transfection of degraders with mCherry providing a measure of transfection efficiency. Degradation is detected by increase in population of quadrant Q1 (mCherry positive, low GFP). **(c)** Workflow: (i) Initial assessment of target degradability using a panel of chimeric E3s in which the substrate-binding domain is replaced by an anti-GFP nanobody (αGFP) as the target-binding motif; (ii) Replace E3s with degrons for E3s or for motifs for targeting directly to the proteasome or for recruiting the autophagy-lysosome machinery; (iii) Replace αGFP with motifs that bind BCL-x_L_ target; (iv) Use these TBM and DM in combination; (v) Amplify degrader potency using coiled-coil oligomerisation domains. **(d)** Flow cytometry assay for measuring degradation of endogenous target using fluorescent labelled anti-target antibody.

We use the target BCL-x_L_ to illustrate the power of a biologics approach to map degradation-target space. BCL-x_L_ is a member of the B-cell lymphoma 2 (BCL-2) family of proteins that regulate apoptosis *via* interactions between pro-and anti-apoptotic members. Consequently, they are the focus of intensive drug discovery efforts ^32^. Although small-molecule inhibitors and PROTACs have been developed for BCL-2 proteins ^33–37^, the need remains for more-potent molecules that have fewer off-target effects. First, we assess the degradability of BCL-x_L_ by testing a panel of chimeric E3s in a cell line that stably expresses a GFP-BCL-x_L_ fusion (**Fig. 1b**). Next, to access the broader cellular landscape of quality-control pathways, we design a wider panel of degradation motifs (DMs in **Fig. 1**) from known degrons for E3s and SLiMs that directly harness the proteasome or the autophagy-lysosome machinery. We combine these degradation motifs with BCL-x_L_-binding motifs (target-binding motifs, TBMs) to make the bifunctional units (**Fig. 1c**).

Specifically, the DMs and TMBs are appended to *de novo* designed coiled-coil sequences that serve as scaffolds and oligomerisation domains for multi-valent presentation of the motifs in diverse configurations (**Fig. 1c(v)**). In total, the protomers are <80 residues long, and they show enhanced potency relative to single-valency counterparts in driving degradation of BCL-x_L_ and the initiation of apoptosis. In so doing, we generate a series of adaptable molecular glues ^38, 39^ that could be used for various applications.

## Results

### GFP-BCL-x_**L**_ is directed for degradation by many different chimeric E3s

We used a stepwise workflow for degrader design (**Fig. 1c**), starting with assessment of degradability of a GFP-BCL-x_L_ fusion protein, followed by testing BCL-x_L_-binding and degradation motifs, then fully engineered hetero-bifunctional molecules, and lastly oligomeric variants to increase the valency assembled using the coiled-coil scaffold ^40^. To explore the degradability of BCL-x_L_, we made stable HEK293T cell lines constitutively expressing full-length BCL-x_L_ fused at its *N* terminus to eGFP (a GFP variant with enhanced fluorescence properties, referred to subsequently as GFP). These cell lines were transfected with a panel of chimeric E3s, in which the natural substrate-binding domain of the E3 substrate-recognition subunit was replaced by the nanobody vhhGFP4 (referred to subsequently as αGFP) that binds with nanomolar affinity to GFP (**Fig. 1c(i), Table S1** and **Table S2**), followed by a triple HA tag, a T2A cleavage site, and mCherry to assess transfection efficiency. The chosen E3s were well-characterised in their degron interactions and had previously been shown to be effective in targeted protein degradation ^40^. Degradation of GFP-BCL-x_L_ was assessed using flow cytometry. Events were gated based on fluorescence of cells stably expressing GFP-BCL-x_L_ and transfected with mCherry-only control (quadrant Q2). Successful degradation of BCL-x_L_ resulted in a decrease in GFP signal intensity and a shift from Q2 to Q1 (**Fig. 2a**). Data are presented as percentages of a remaining GFP-BCL-x_L_: (1-Q1/(Q1+Q2))*100.

**Fig. 2.**
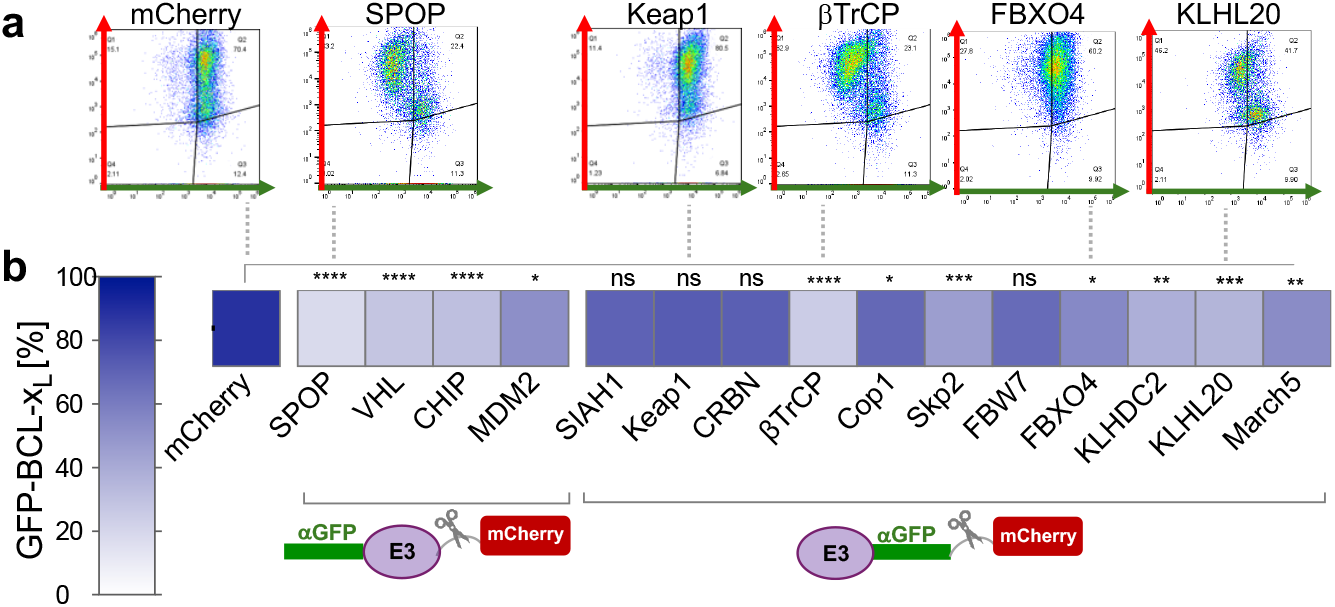
GFP-Bcl-x_L_ is degraded by αGFP-based chimeric E3s. Flow cytometry analysis of HEK293T cell line constitutively expressing GFP-Bcl-xL 48 hours post transient transfection with the chimeric E3s, in which the natural substrate-binding domain of the E3 or E3 substrate-recognition subunit was replaced by αGFP. The schematics at the bottom of the figure indicate whether αGFP is at the N- or C-terminus of the construct. mCherry was used as a negative control (see also **Fig. S1B**). Statistics were calculated using Welsch’s t test.

We generated two cell lines, one with GFP-BCL-x_L_ expression under the weak HSV TK (herpes simplex virus thymidine kinase) promoter, and one under the strong CMV (cytomegalovirus) promoter. As expected, the former expressed lower GFP-BCL-x_L_ protein levels than the latter (**Fig. S1a**). Upon transient transfection with the chimeric E3s, similar effects on GFP-BCL-x_L_ protein levels were observed in the two cell lines. However, the much higher BCL-x_L_ expression resulting from the stronger promoter led to overall lower extents of degradation, and consequently the BCL-x_L_ construct with the weaker promoter was used for subsequent experiments (**Fig. S1b**). The following E3s were found to support degradation of BCL-x_L_: SPOP, VHL, CHIP, βTrCP, KLHDC2, and KLHL20 (**Fig. 2**). We confirmed expression of the chimeric E3s by Western blot (**Fig. S2A**).

### Degraders can be designed to harness other degradation machineries

Next, we sought to co-opt other cellular quality-control pathways for BCL-x_L_ degradation by making constructs containing one or multiple copies of different degradation motifs fused to αGFP (Fig. 1c(ii) and Table S1). The following degradation motifs were used: degrons from natural substrates of those E3s for which the chimeras were found to induce GFP-BCL-x_L_ degradation (namely SPOP, VHL, CHIP, βTrCP, KLHDC2, KLHL20), and a degron recognised by the UBR family of E3s ^41–47^; a PEST sequence (found in short-lived proteins) from the protein ornithine decarboxylase (ODC) ^48^; a short peptide sequence (known as MC2) recently identified from a library screen to bind to the proteasome ^24^; 1 – 3 copies of UBL domain(s) from proteasome “shuttle” proteins fused to αGFP ^25, 49^; LC3-interaction regions (LIRs) to recruit the LC3/GABARAP family of autophagy receptors ^50, 51^. We included the chimeric SPOP E3 as a positive control. The data showed that all pathways can be co-opted by these degradation motifs to drive the destruction of GFP-BCL-x_L_ (**Fig. 3**). Degrons for the E3s SPOP, VHL, KLHL20, and UBR, MC2 and (UBL)_3_ for direct targeting to the proteasome, and LIR2 for autophagy targeting resulted in the most degradation (40 – 50% relative to the mCherry control), and they were as potent as the chimeric SPOP E3 (see also **Fig. S3**). Interestingly, the degron for UBR induced degradation when placed at the *C* terminus of αGFP, even though this is known as an *N-*terminal degron. It is possible that this peptide acts as a *C-*terminal destabilisation domain, as has been observed previously ^52^. For the proteasome-mediated degraders, co-treatment with proteasome inhibitor Carfilzomib resulted in less GFP-BCL-x_L_ degradation, although the effects were variable (**Fig. S4A**).

**Fig. 3.**
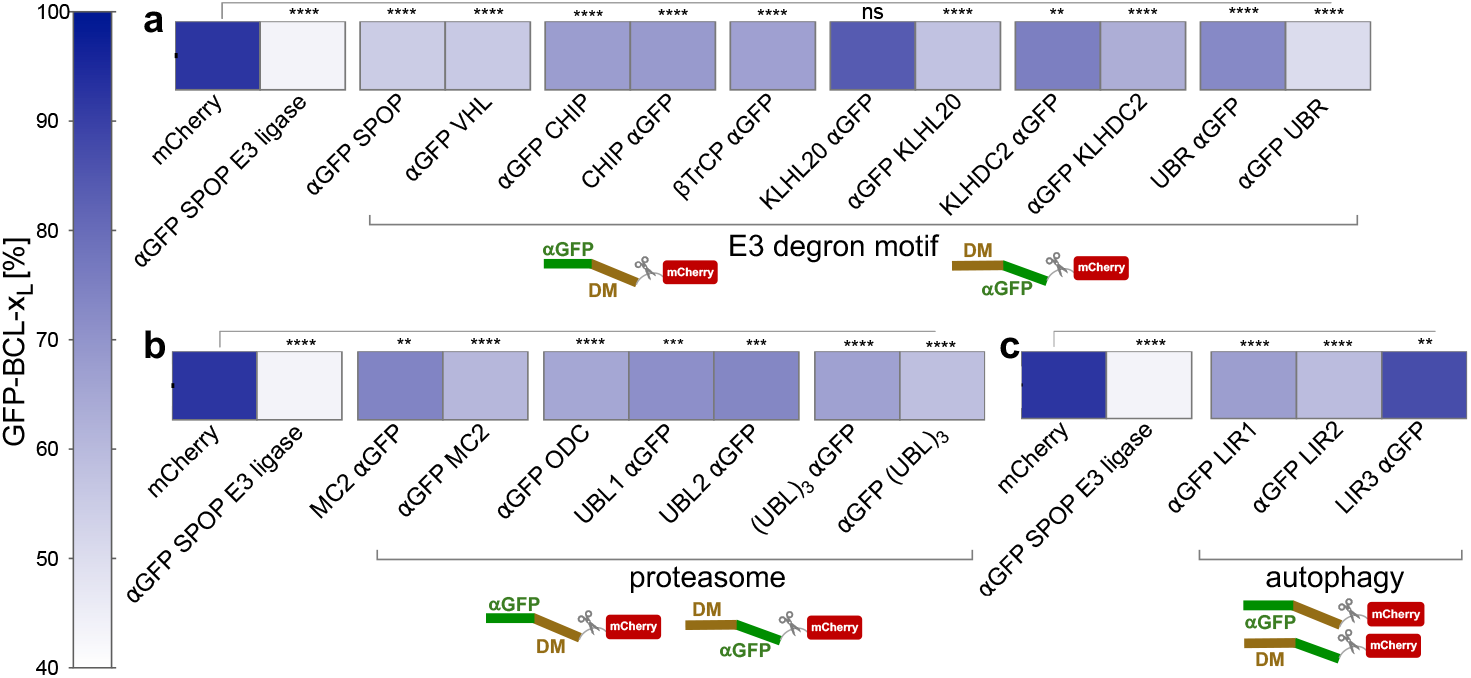
GFP-BCL-x_L_ is degraded by αGFP-degron fusions. Flow cytometry analysis of HEK293T cells constitutively expressing GFP-BCL-x_L_ 48 hours post transfection with **(a)** degraders composed of degrons for E3s, **(b)** degrons targeting directly to the proteasome, and **(c)** degrons targeting to the autophagy-lysosome pathway (see also **Fig. S3 for bar plots**). Statistics were calculated using Welsch’s t test.

### Combining BCL-x_L_-binding and degradation motifs

Next, we replaced the αGFP nanobody with a panel of both natural and synthetic BCL-x_L_ binders: short (19 – 25 residue) peptides from the BH3 proteins BAD and HRK ^53, 54^; a BAD-HRK chimeric peptide optimized for BCL-x_L_ selectivity (referred to here as BAD-HRK); and an adhiron (a synthetic single-domain antibody-like molecule, referred to here as Ad_BCL-xL_) identified previously by members of our team (referred to in our earlier work as Af7 - Affimer7) ^55^. These binders all have dissociation constants in the nanomolar range, albeit lower than the affinity of the αGFP nanobody for GFP (**Table S2**). We fused these BCL-x_L_ binders to the following degradation motifs that we found effective in combination with αGFP (**Fig. 3**); namely, the UBR degron, and the ODC and MC2 peptides for direct targeting to the proteasome. We tested these degraders for binding to GFP-BCL-x_L_. For comparison, we included the αGFP-based degraders, and we also made BCL-x_L_ binder-SPOP chimeras. The data (**Fig. 4a**) revealed that BAD-UBR, BAD-MC2, and HRK-UBR were as potent, or more potent than the αGFP-based degraders and the SPOP chimeras. BCL-x_L_ degradation and degrader expression were confirmed by Western blot (**Fig. S5**). Two timepoints, 24 h and 48 h, were taken, with more BCL-x_L_ degradation being observed for the latter. Degrader levels remained relatively constant over this timescale.

**Fig. 4.**
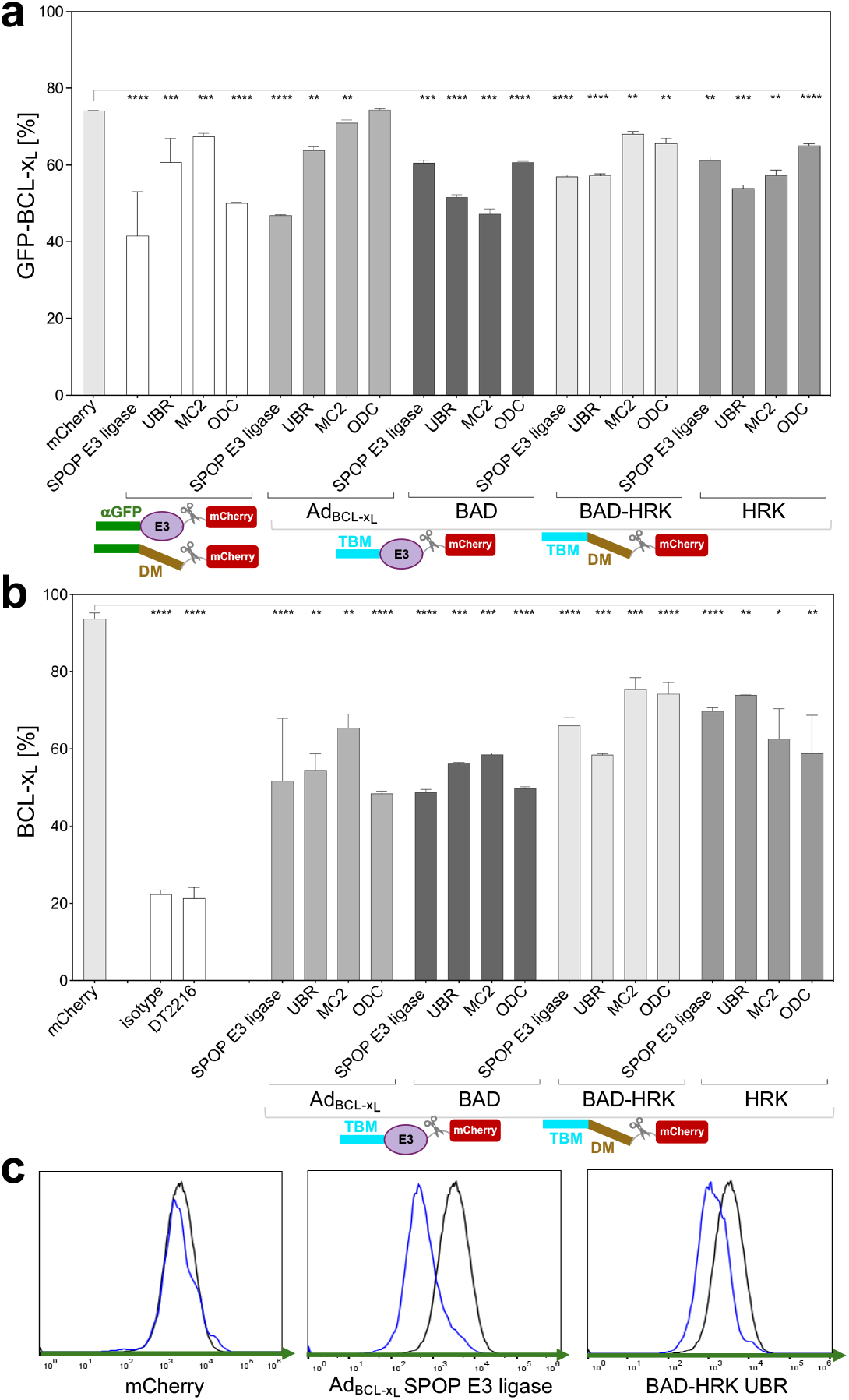
Degradation harnessing high-affinity BCL-x_L_ binders. **(a)** Flow cytometry analysis of HEK293T cell line constitutively expressing GFP-BCL-x_L_ 48 hours post transfection with the degraders. mCherry-only was used as a negative control. αGFP-based degraders were tested in parallel. Statistics were calculated using Welsch’s t test. **(b)** Flow cytometry analysis of endogenous BCL-x_L_ levels in A549 cell line 48 hours post transfection with degraders. Isotype refers to IgG control. mCherry-only was used as a negative control. See **Fig. S6** for additional control molecules **(c)** Representative histograms of the flow cytometry data showing degrader in blue and the mCherry control in black.

### Time-dependence of degradation of Nanoluc-BCL-x_**L**_

To confirm the above results, and to explore the time-course of degradation, a cell line stably expressing Nanoluc-BCL-x_L_ was established and degradation was measured by luminescence over a 24-hour period post transfection with the chimeric degraders or αGFP-only control. This showed an initial increase in luminescence over 2 hours, which arises from the luciferase substrate being converted to its active form in the cell (**Fig. S7**). After ∼4 hours the luminescence decreased indicating degradation of BCL-x_L_. No decrease in luminescence was observed for the αGFP-only control.

### Degraders reduce levels of endogenous BCL-x_**L**_

Next, we selected the most potent degraders of GFP-BCL-x_L_ and tested them against endogenous BCL-x_L_. We used the A549 cell line, which expresses BCL-x_L_ and is BCL-2 family-dependent (treatment with inhibitor ABT-263 (Navitoclax) combined with siRNA for MCL-1 suppression induces apoptosis) ^56^. Flow cytometry with a fluorescently labelled anti-BCL-x_L_ antibody was used to monitor BCL-x_L_ levels. The data indicate that endogenous BCL-x_L_ levels reduced upon degrader treatment, with the Ad_BCL-xL_- and BAD-based degraders being slightly more potent than the HRK- and BAD-HRK-based degraders (**Fig. 4b**). Controls, including SPOP chimeras comprising an Ad_BCL-xL_ mutant that abrogates BCL-x_L_ binding, and a peptide that bind to other BCL-2 family members but not to BCL-x_L_ (BAD mutant), did not induce degradation (**Fig. S6**). The BCL inhibitor Navitoclax also had no effect on BCL-x_L_ levels, whereas the small-molecule BCL-x_L_ PROTAC DT2216 (comprising Navitoclax and a VHL ligand connected via a linker ^37^) did induce degradation. Again, degrader expression was confirmed by Western blot (**Fig. S5)**.

### BCL-x_**L**_ degraders induce apoptosis

We tested the degraders for their ability to induce apoptosis, and for cell viability and cytotoxicity. We used a flow-cytometry assay, in which binding of FITC-labelled Annexin V to externalised phosphatidylserine (PS) provides a measure of the early stages of apoptosis. In addition, cells in late stages of apoptosis lose membrane integrity and stain with a viability dye such as DAPI. Apoptosis was calculated as the sum of cells in early and late stages expressed as percentage of total cell count. The degraders had only a small effect on cell viability and exhibited some cytotoxic effect (**Fig. S8**), whereas they induced significant levels of apoptosis (**Fig. 5**). The BAD-based and BAD-HRK-based degraders were more potent than Ad_BCL-xL_- and HRK-based degraders in induction/stimulation of apoptosis, with several observed to be as potent as the BCL-x_L_ PROTAC DT2216. Staurosporine, a standard apoptosis inducer, showed non-specific cytotoxicity.

**Fig. 5.**
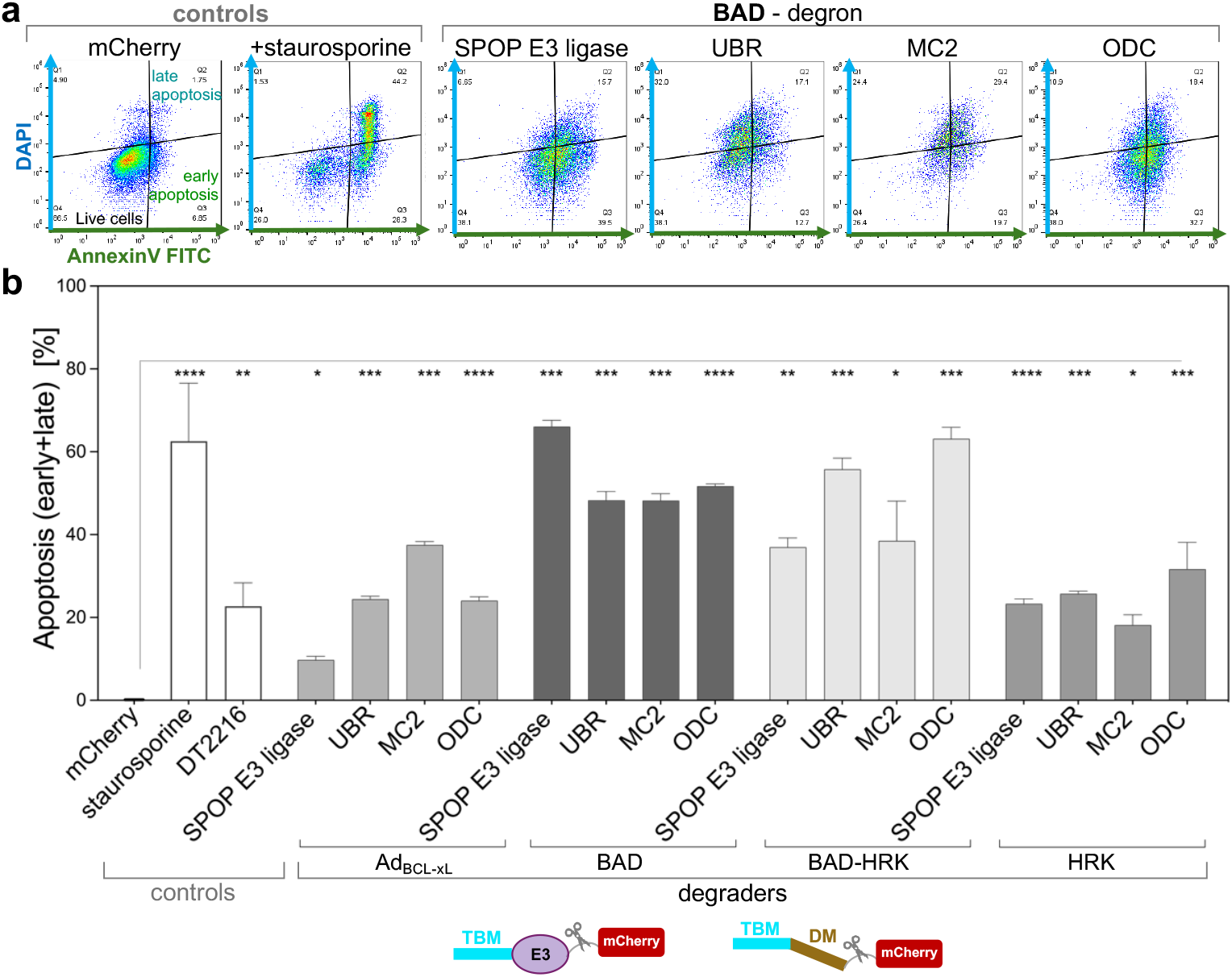
Induction of apoptosis upon transfection with BCL-x_L_ degraders. Measurements were made 48 hours post transfection with the degraders.

### Coiled-coil scaffolds bring multi-valency to degrader design enhancing potency

Lastly, we explored the oligomerisation of the coiled-coil (CC) scaffolds to increase the valency of the degraders. To this end, we appended BCL-x_L_-binding peptides (target-binding motifs, TBM) and degradation motifs (DM) onto one or both termini of different *de novo* designed coiled-coil oligomerisation sequences (**Fig. 1c(v)**). We assembled these degraders in a combinatorial fashion from the following motifs linked by poly(GS) spacers (Table S1): (i) the BAD peptide as the BCL-x_L_ binding motifs; (ii) two different degrons (UBR and ODC); and (iii) four different *de novo* designed 30-residue coiled-coil modules (anti-parallel dimer, parallel trimer, parallel and anti-parallel tetramer) ^28, 47, 48, 54, 57, 58^. As shown in **Fig. 6a**, we tested different arrangements of TBM and DM alongside single-valency analogues, and we compared them with the original degrader analogues that contained simple linear concatenations of the degrons. The data (**Fig. 6b and Fig. S9**) revealed that in general the potency of the degraders increased with increasing valency.

**Fig. 6.**
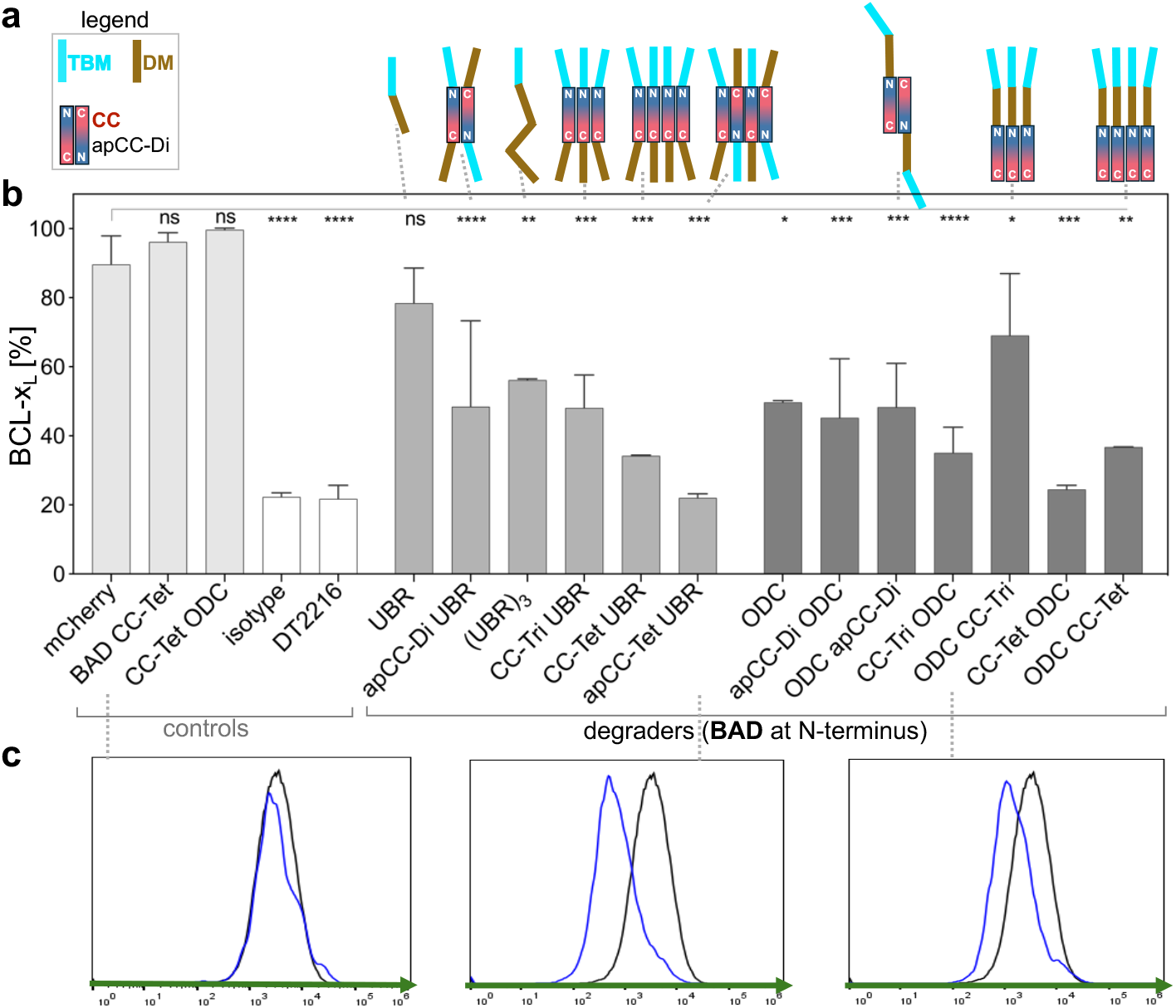
BCL-x_L_ degradation and induction of apoptosis using multivalent degraders. **(a)** Schematic showing coiled-coil (CC) designs and, for comparison, single or three tandem repeats of the degradation motif; **(b)** and **(c)** Degradation of endogenous BCL-x_L_ analysed by flow cytometry 48 hours post transient transfection with the degraders. Data for additional degrader designs are shown in **Fig. S9**.

Moreover, the potency varied depending on the configuration of BCL-x_L_ binder and degron (e.g. BAD-CC-Tri-ODC versus BAD-ODC CC-Tri). Thus, we can encode target-binding and effector functions into small (∼80-residue) coiled-coil-based sequences, which are as potent degraders as the much larger chimeric E3s (e.g. the Ad_BCL-xL_-SPOP chimera is 344 residues). They are also as potent as the small molecule PROTAC DT2216 ^59^. A subset of degraders was further tested in the apoptosis assay, which confirmed that degradation of BCL-x_L_ elicits apoptosis (**Fig. S10**).

## Discussion

TPD is an exciting therapeutic modality with tremendous potential. However, it is constrained by the lack of small-molecule ligands to both therapeutic targets and much of the cell’s quality-control machineries, including most of the hundreds of E3 ubiquitin ligases in the human proteome. Here we introduce a biologics approach to expand the target and degradation space using the anti-apoptotic target BCL-x_L_ as a model target. First, we use chimeric E3s to explore the degradability of BCL-x_L_. Next, we show that the degradation landscape can be expanded by using SLiMs to bind BCL-x_L_ target (target-binding motifs, TBM) combined with SLiMs to recruit either E3s, the proteasome, or the autophagy-lysosome machinery (degradation motifs, DM). In order to (1) integrate TBM and DM into a single entity, (2) enhance the potency of these motifs, and (3) diversify their geometries, we take advantage of versatile *de novo* coiled-coil scaffolds. The second of these three goals is inspired by the frequent use in nature of multivalency to increase binding affinities and specificity. In the ubiquitin-proteasome system, multivalent recognition mechanisms may facilitate optimal substrate positioning on the E3 and processivity of ubiquitination.^60^ To achieve multivalency within synthetic degraders, we construct coiled-coil dimers, trimers, and tetramers appended with the BCL-x_L_-binding and degradation motifs. We show that they were more effective than their single-valency counterparts in driving BCL-x_L_ degradation and inducing apoptosis, and the higher the valency (dimeric *versus* trimeric *versus* tetrameric coiled coils) the more effective they are.

In summary, the 80-residue hetero-bifunctional coiled coils developed here illustrate the efficiency with which this architecture can be harnessed to construct a simple plug-and-play TPD platform that enables mapping and optimising degradation-target space without the need for sophisticated design methodologies. We call this the CoCoProx (<u>Co</u>iled <u>Co</u>ils for induced <u>Prox</u>imity) platform and propose that it could be used to drive degradation of many currently undrugged targets as well as for other induced-proximity reactions.

## Materials and Methods

### Generation of cell lines stably expressing GFP- and Nanoluc-tagged BCL-x_**L**_

All cell lines were incubated at 37 °C with 5% CO_2_, 95% air and ≥95% humidity. HEK293T cells were grown in Dulbecco’s Modified Eagle Medium (DMEM) (GibcoTM) supplemented with 10% FBS (fetal bovine serum), and A549 cells were grown in Nutrient Mixture F-12 Ham medium (Merck) supplemented with 10% FBS. DNA transfections were performed using cationic lipids TransIT-2020 (Mirus Bio). A stable cell line constitutively expressing Nanoluc-BCL-x_L_ was generated by an antibiotic selection to isolate cells that successfully incorporated the genetic material. Stable GFP-BCL-x_L_ cell lines were generated by antibiotic selection pressure prior to sorting of live GFP-positive cells for further propagation (using the BD FACS Aria instrument in the Flow Cytometry facility, Department of Pathology, University of Cambridge).

### Molecular biology

Chimeric E3s containing an αGFP nanobody with a 3xHA tag at the N-terminus and under a doxycycline-inducible promoter were constructed in plasmid pcDNA5 FRT/TO (see SI Table S1 for further details). Gene fragments encoding other degraders were ordered as gBlocks (IDT) and cloned into the plasmid pcDNA5 FRT/TO using NEBuilder^®^ HiFi DNA Assembly Master Mix (NEB) according to the manufacturer’s instructions. These degraders also have 3xHA N-terminal tag and are expressed under a doxycycline-inducible promoter. Gene fragments encoding the multivalent degraders were synthesised in the pcDNA3.1 Hygro(+) plasmid by Genscript; these degraders have a FLAG tag at the N-terminus and are constitutively expressed. All degraders have a C-terminal mCherry preceded by a T2A self-cleaving peptide sequence.

### Flow cytometry assay of degradation of GFP-BCL-x_**L**_ **and endogenous BCL-x**_**L**_

Cells were seeded in 24-well plates at a density of 300,000 cells in supplemented DMEM without antibiotics. The following morning, transfection of chimeras was performed using Mirus TransIT-2020 as per manufacturer’s protocol, using 0.5 μg DNA: 1.5 μl transfection reagent. 24 hours post-transfection doxycycline at a concentration of 100 ng/ml was added, and 48 hours after transfection, wells were washed with PBS, trypsinized, resuspended in FACS buffer (PBS with 2% FBS), and stained with 1 μg/ml DAPI, prior to flow cytometry analysis using an Attune NxT flow cytometer (ThermoFisher Scientific). All analysis was performed using the FlowJo V10 software. Events were gated based on fluorescence of cells stably expressing GFP-BCL-x_L_ and transfected with mCherry-only control (quadrant Q2). Successful degradation of BCL-x_L_ resulted in a decrease in GFP signal intensity and a shift to Q1. Data are from at least 3 independent experiments and shown as percentages of a remaining GFP-BCL-x_L_ (1-Q1/(Q1+Q2))*100 ± standard deviation (SD). Statistics were calculated using Welsch’s t test. Not significant (ns) ⩾ 0.05; \**P* < 0.05; \*\**P* < 0.01; \*\*\**P* < 0.001; \*\*\*\**P* < 0.0001.

For detection of endogenous BCL-x_L_, cells were collected, washed, and fixed in 4% formaldehyde. Then, following wash with PBS cells were permeabilized with cold methanol for at least 15 min on ice and incubated overnight at 4°C with BCL-x_L_ (54H6) Rabbit mAb (Alexa Fluor 488 Conjugate) antibody and a Rabbit IgG Isotype control, Alexa Fluor 488 Conjugated (Cell Signaling, BS-0295P-A488-BSS-100 μl Stratech Scientific Ltd, respectively). Subsequently, cells were washed with PBS and resuspended in PBS for FACS analysis. Results were plotted as events of BCL-x_L_ Alexa 488 channel. The percentage of cells that lost GFP fluorescence was calculated by subtracting out the florescence of the control (mCherry) using the Overton positive method using the FlowJo V10 software. Data are from at least 3 independent experiments. Values are presented as the mean ± SD. Statistics were calculated using Welsch’s t test. ns ⩾ 0.05; \**P* < 0.05; \*\**P* < 0.01; \*\*\**P* < 0.001; \*\*\*\**P* < 0.0001.

### Luminescence-based assay for time dependence of Nanoluc-tagged BCL-x_**L**_ degradation

HEK293T cell lines stably expressing Nanoluc-BCL-x_L_ were seeded and used at approximately 70% confluency. Cells were transfected with degraders, and 24 hours post transfection 1/1000 endurazine (Promega) was added. Luminescence was measured with a 470-80 nm filter at 5-minute intervals using a CLARIOstar Plus plate reader (BMG) at 37 °C, 5% CO_2_, 95% air.

### Western blot

Cell cultures to be harvested were washed and lysed according to standard procedures for western blotting. All protein transfers were performed via semi-dry electroblotting onto PVDF membranes (Immobilin-P, Merck) using a Pierce Power Blot Cassette (ThemoFisher Scientific). Transfers were typically performed with a voltage of 15 V and a current of 2.5 A for 30 minutes. Membranes were cut, as required, to allow separate staining of proteins of different masses and blocked for 1 hour in 5% milk (Marvel) at room temperature with gentle agitation. Primary antibody stains were typically performed overnight at 4 °C in 5% milk. The following morning, stained membranes were washed three times for 5 minutes with PBS-T (PBS, 0.1% Tween 20). HRP-conjugated secondary antibody stains were performed according to primary antibody species at a concentration of 1/10,000. Staining with Rabbit Anti-BCL-x_L_-HRP antibody (Cell Signaling) and mouse anti-Nanoluc-HRP monoclonal antibody (Promega) was performed according to the manufacturer’s instructions. PBS-T washes were repeated as above before imaging. All HRP-stained membranes were developed using Amersham ECL Western Blotting Detection Reagent (Cytiva). Detection of chemiluminescence was performed using an Odyssey Fc Imaging System (LI-COR).

### Apoptosis assay

Viability and cytotoxicity assays were performed using Promega kits according to their protocols, with the exception of additional filters that we used to eliminate the background fluorescence from the mCherry co-expressed with degraders; fluorescence filter wavelengths were 485±10 nm for excitation and 520±10nm for emission for the cytotoxicity measurements, and 400±20 nm for excitation and 505±20 nm for emission for the viability measurements. Induction of apoptosis was analysed using flow cytometry of FITC-labelled Annexin V and DAPI staining (BioLegend and BD Pharmingen, respectively). Briefly, 48 hours post-transfection of A549 cells with degraders, PROTAC, or staurosporine control, cells were detached, collected in tubes, washed with PBS and incubated for 30 min to recover. Subsequently, cells were stained with FITC-conjugated Annexin V in accordance with the manufacture’s protocol except for a longer incubation time of 45 min was used. Next, cells were washed in 1x binding buffer and resuspended in a 300 μl of this buffer containing a viability dye (DAPI). Flow cytometry measurements were performed within 2 hours using the appropriate fluorophore channels for FITC and DAPI. Cells undergoing early apoptosis have externalized phosphatidylserine (PS) that binds to Annexin in a calcium-dependent manner. Cells in late stages of apoptosis lose membrane integrity and stain with a viability dye such as DAPI. Apoptosis was calculated as the sum of cells in early and late stages and expressed as percentages of total cell count.

## Acknowledgements

This work was supported by BBSRC grants BB/V008412/1 and BB/V008412/2 to AJW, LSI, DNW, and TAE, BB/Y007816/1 to LSI, and BB/V003577/1 and BB/V003577/2 to AJW. D.T.H. was supported by Cancer Research UK (A29256).

