## Supplemental Information for "Coiled-coil scaffolds for Targeted Protein Degradation"

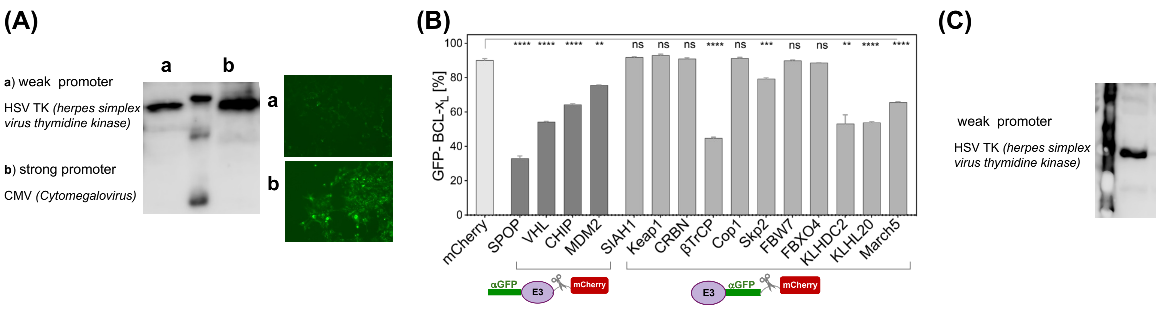


**Figure S1. (A)** Western blot (left) and microscopy (right) of engineered HEK293T cell lines constitutively expressing GFP-BCL-$x_L$ on a weak promoter HSV TK (a) and a strong promoter CMV (b) probed with anti-GFP-HRP antibody (#GTX628528-01, Insight Biotechnology). **(B)** Degradation of GPF-BCL-$x_L$ expressed under the strong promoter following transient transfection of chimeric E3s. The schematics at the bottom of the figure indicate whether αGFP is positioned at the N- or C-terminus of the construct. mCherry served as a negative control. Results are from at least three independent experiments and presented as the mean ± SD. Statistics were calculated using Welsch's t test (ns ⩾ 0.05; ****P* < 0.01; *****P* < 0.001; ******P* < 0.0001).
**(C)** Western blot of engineered HEK293T cell lines constitutively expressing Nanoluc-BCL-$x_L$ with Anti-NanoLuc® Monoclonal Antibody (Promega #N7000).

**Table S1. Amino acid sequences of constructs.**

| | |
|---|---|
| **Constructs for stable cell lines generation** | |
| eGFP-BCL-$x_L$ | MVSKGEELFTGVVPILVELDGDVNGHKFSVSGEGEGDATYGKLTLKFICTTGKLPVPWPTLVTTLTYGVQCFSRYPDHMKQHDFFKSAMPEGYVQERTIFFKDDGNYKTRAEVKFEGDTLVNRIELKGIDFKEDGNILGHKLEYNYNSHNVYIMADKQKNGIKVNFKIRHNIEDGSVQLADHYQQNTPIGDGPVLLPDNHYLSTQSALSKDPNEKRDHMVLLEFVTAAGITLGMDELYKVDPVAGAggsggggsMSQSNRELVVDFLSYKLSQKGYSWSQFSDVEENRTEAPEGTESEMETPSAINGNPSWHLADSPAVNGATAHSSSLDAREVIPMAAVKQALREAGDEFELRYRRAFSDLTSQLHITPGTAYQSFEQVVNELFRDGVNWGRIVAFFSFGGALCVESVDKEMQVLVSRIAAWMATYLNDHLEPWIQENGGWDTFVELYGNNAAAESRKGQERFNRWFLTGMTVAGVVLLGSLFSRK* |
| Nanoluc-BCL-$x_L$ | MVFTLEDFVGDWRQTAGYNLDQVLEQGGVSSLFQNLGVSVTPIQRIVLSGENGLKIDIHVIIPYEGLSGDQMGQIEKIFKVVYPVDDHHFKVILHYGTLVIDGVTPNMIDYFGRPYEGIAVFDGKKITVTGTLWNGNKIIDERLINPDGSLLFRVTINGVTGWRLCERILAgssggsggggsMSQSNRELVVDFLSYKLSQKGYSWSQFSDVEENRTEAPEGTESEMETPSAINGNPSWHLADSPAVNGATAHSSSLDAREVIPMAAVKQALREAGDEFELRYRRAFSDLTSQLHITPGTAYQSFEQVVNELFRDGVNWGRIVAFFSFGGALCVESVDKEMQVLVSRIAAWMATYLNDHLEPWIQENGGWDTFVELYGNNAAAESRKGQERFNRWFLTGMTVAGVVLLGSLFSRK* |
| **Chimeric E3 ligases** (αGFP located at N- or C-terminus in the same position as the native substate-binding domain)[1] | |
| 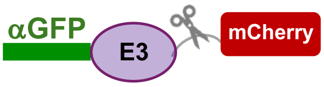  | |
| 3xHA-**αGFP-E3_ligase-**T2A-mCherry | YPYDVPDYAYPYDVPDYAYPYDVPDYAgsdqVQLVESGGALVQPGGSLRLSCAASGFPVNRYSMRWYRQAPGKEREWVAGMSSAGDRSSYEDSVKGRFTISRDDARNTVYLQMNSLKPEDTAVYYCNVNVGFEYWGQGTQVTVSSgssgsg(**E3_ligase**)gssggsgEGRGSLLTCGDVEENPG↑PggsgVSKGEEDNMAIIKEFMRFKVHMEGSVNGHEFEIEGEGEGRPYEGTQTAKLKVTKGGPLPFAWDILSPQFMYGSKAYVKHPADIPDYLKLSFPEGFKWERVMNFEDGGVVTVTQDSSLQDGEFIYKVKLRGTNFPSDGPVMQKKTMGWEASSERMYPEDGALKGEIKQRLKLKDGGHYDAEVKTTYKAKKPVQLPGAYNVNIKLDITSHNEDYTIVEQYERAEGRHSTGGMDELYK* |
| 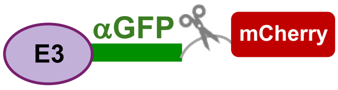  | |
| 3xHA- **E3_ligase-αGFP-**T2A-mCherry | YPYDVPDYAYPYDVPDYAYPYDVPDYAgs(**E3_ligase**)gssgsgdqVQLVESGGALVQPGGSLRLSCAASGFPVNRYSMRWYRQAPGKEREWVAGMSSAGDRSSYEDSVKGRFTISRDDARNTVYLQMNSLKPEDTAVYYCNVNVGFEYWGQGTQVTVSSgssggsgEGRGSLLTCGDVEENPG↑PggsgVSKGEEDNMAIIKEFMRFKVHMEGSVNGHEFEIEGEGEGRPYEGTQTAKLKVTKGGPLPFAWDILSPQFMYGSKAYVKHPADIPDYLKLSFPEGFKWERVMNFEDGGVVTVTQDSSLQDGEFIYKVKLRGTNFPSDGPVMQKKTMGWEASSERMYPEDGALKGEIKQRLKLKDGGHYDAEVKTTYKAKKPVQLPGAYNVNIKLDITSHNEDYTIVEQYERAEGRHSTGGMDELYK* |

| E3 ligases[1] | |
|---|---|
| 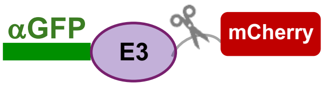 | |
| SPOP$_{167-374}$ | SVNISGQNTMNMVKVPECRLADELGGLWENSRFTDCCLCVAGQEFQAHKAILAARSPVFSAMFEHEMEESKKNRVEINDVEPEVFKEMMCFIYTGKAPNLDKMADDLLAAADKYALERLKVMCEDALCSNLSVENAAEILILADLHSADQLKTQAVDFINYHASDVLETSGWKSMVVSHPHLVAEAYRSLASAQCPFLGPPRKRLKQS |
| VHL$_{152-213}$ | TLPVYTLKERCLQVVRSLVKPENYRRLDIVRSLYEDLEDHPNVQKDLERLTQERIAHQRMGD |
| CHIP$_{128-303}$ | RLNFGDDIPSALRIAKKKRWNSIEERRIHQESELHSYLSRLIAAERERELEECQRNHEGDEDDSHVRAQQACIEAKHDKYMADMDELFSQVDEKRKKRDIPDYLCGKISFELMREPCITPSGITYDRKDIEEHLQRVGHFDPVTRSPLTQEQLIPNLAMKEVIDAFISENGWVEDY |
| MDM2$_{111-491}$ | NQQESSDSGTSVSENRCHLEGGSDQKDLVQELQEEKPSSSHLVSRPSTSSRRRAISETEENSDELSGERQRKRHKSDSISLSFDESLALCVIREICCERSSSSESTGTPSNPDLDAGVSEHSGDWLDQDSVSDQFSVEFEVESLDSEDYSLSEEGQELSDEDDEVYQVTVYQAGESDTDSFEEDPEISLADYWKCTSCNEMNPPLPSHCNRCWALRENWLPEDKGKDKGEISEKAKLENSTQAEEGFDVPDCKKTIVNDSRESCVEENDDKITQASQSQESEDYSQPSTSSSIIYSSQEDVKEFEREETQDKEESVESSLPLNAIEPCVICQGRPKNGCIVHGKTGHLMACFTCAKKLKKRNKPCPVCRQPIQMIVLTYFP |
| 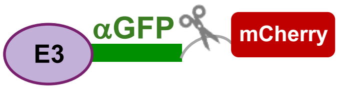 | |
| SIAH1$_{2-89}$ | SRQTATALPTGTSKCPPSQRVPALTGTTASNNDLASLFECPVCFDYVLPPILQCQSGHLVCSNCRPKLTCCPTCRGPLGSIRNLAMEK |
| Keap1$_{2-314}$ | QPDPRPSGAGACCRFLPLQSQCPEGAGDAVMYASTECKAEVTPSQHGNRTFSYTLEDHTKQAFGIMNELRLSQQLCDVTLQVKYQDAPAAQFMAHKVVLASSSPVFKAMFTNGLREQGMEVVSIEGIHPKVMERLIEFAYTASISMGEKCVLHVMNGAVMYQIDSVVRACSDFLVQQLDPSNAIGIANFAEQIGCVELHQRAREYIYMHFGEVAKQEEFFNLSHCQLVTLISRDDLNVRCESEVFHACINWVKYDCEQRRFYVQALLRAVRCHSLTPNFLQMQLQKCEILQSDSRCKDYLVKIFEELTLHKPT |
| CRBN$_{2-320}$ | AGEGDQQDAAHNMGNHLPLLPAESEEEDEMEVEDQDSKEAKKPNIINFDTSLPTSHTYLGADMEEFHGRTLHDDDSCQVIPVLPQVMMILIPGQTLPLQLFHPQEVSMVRNLIQKDRTFAVLAYSNVQEREAQFGTTAEIYAYREEQDFGIEIVKVKAIGRQRFKVLELRTQSDGIQQAKVQILPECVLPSTMSAVQLESLNKCQIFPSKPVSREDQCSYKWWQKYQKRKFHCANLTSWPRWLYSLYDAETLMDRIKKQLREWDENLKDDSLPSNPIDFSYRVAACLPIDDVLRIQLLKIGSAIQRLRCELDIMNKCTS |
| βTrCP$_{2-263}$ | DPAEAVLQEKALKFMNSSEREDCNNGEPPRKIIPEKNSLRQTYNSCARLCLNQETVCLASTAMKTENCVAKTKLANGTSSMIVPKQRKLSASYEKEKELCVKYFEQWSESDQVEFVEHLISQMCHYQHGHINSYLKPMLQRDFITALPARGLDHIAENILSYLDAKSLCAAELVCKEWYRVTSDGMLWKKLIERMVRTDSLWRGLAERRGWGQYLFKNKPPDGNAPPNSFYRALYPKIIQDIETIESNWR |
| Cop1$_{2-836}$ | SGSRQAGSGSAGTSPGSSAASSVTSASSSLSSSPSPPSVAVSAAALVSGGVAQAAGSGGLGGPVRPVLVAPAVSGSGGGAVSTGLSRHSCAARPSAGVGGSSSSLGSGSRKRPLLAPLCNGLINSYEDKSNDFVCPICFDMIEEAYMTKCGHSFCYKCIHQSLEDNNRCPKCNYVVDNIDHLYPNFLVNELILKQKQRFEEKRFKLDHSVSSTNGHRWQIFQDWLGTDQDNLDLANVNLMLELLVQKKKQLEAESHAAQLQILMEFLKVARRNKREQLEQIQKELSVLEEDIKRVEEMSGLYSPVSEDSTVPQFEAPSPSHSSIIDSTEYSQPPGFSGSSQTKKQPWYNSTLASRRKRLTAHFEDLEQCYFSTRMSRISDDSRTA |
| Skp2$_{2-150}$ | HRKHLQEIPDLSSNVATSFTWGWDSSKTSELLSGMGVSALEKEEPDSENIPQELLSNLGHPESPPRKRLKSKGSDKDFVIVRRPKLNRENFPGVSWDSLPDELLLGIFSCLCLPELLKVSGVCKRWYRLASDESLWQTLDLTGKNLHPD |

| | |
|---|---|
| FBW7$_{2-293}$ | CVPRSGLILSCICLYCGVLLPVLLPNLPFLTCLSMSTLESVTYLPEKGLYCQRLP<br>SSRTHGGTESLKGKNTENMGFYGTLKMIFYKMKRKLDHGSEVRSFSLGKKPC<br>KVSEYTSTTGLVPCSATPTTFGDLRAANGQGQQRRRITSVQPPTGLQEWLKM<br>FQSWSGPEKLLALDELIDSCEPTQVKHMMQVIEPQFQRDFISLLPKELALYVLS<br>FLEPKDLLQAAQTCRYWRILAEDNLLWREKCKEEGIDEPLHIKRRKVIKPGFIH<br>SPWKSAYIRQHRIDTNWRRGELKSP |
| FBXO4 | MAGSEPRSGTNSPPPPFSDWGRLEAAILSGWKTFWQSVSKERVARTTSREE<br>VDEAASTLTRLPIDVQLYILSFLSPHDLCQLGSTNHYWNETVRDPILWRYFLLR<br>DLPSWSSVDWKSLPDLEILKKPISEVTDGAFFDYMAVYRMCCPYTRRASKSS<br>RPMY |
| KLHDC2 | ADGNEDLRADDLPGPAFESYESMELACPAERSGHVAVSDGRHMFVWGGYK<br>SNQVRGLYDFYLPREELWIYNMETGRWKKINTEGDVPPSMSGSCAVCVDRV<br>LYLFGGHHSRGNTNKFYMLDSRSTDRVLQWERIDCQGIPPSSKDKLGVWVYK<br>NKLIFFGGYGYLPEDKVLGTFEFDETSFWNSSHPRGWNDHVHILDTETFTWS<br>QPITTGKAPSPRAAHACATVGNRGFVFGGRYRDARMNDLHYLNLDTWEWNE<br>LIPQGICPVGRSWHSLTPVSSDHLFLFGGFTTDKQPLSDAWTYCISKNEWIQF<br>NHPYTEKPRLWHTACASDEGEVIVFGGCANNLLVHHRAAHSNEILIFSVQPKS<br>LVRLSLEAVICFKEMLANSWNCLPKHLLHSVNQRFGSNNTSGS |
| KLHL20$_{1-318}$ [2, 3] | MEGKPMRRCTNIRPGETGMDVTSRCTLGDPNKLPEGVPQPARMPYISDKHP<br>RQTLEVINLLRKHRELCDVVLVVGAKKIYAHRVILSACSPYFRAMFTGELAESR<br>QTEVVIRDIDERAMELLIDFAYTSQITVEEGNVQTLLPAACLLQLAEIQEACCEF<br>LKRQLDPSNCLGIRAFADTHSCRELLRIADKFTQHNFQEVMESEEFMLLPANQ<br>LIDIISSDELNVRSEEQVFNAVMAWVKYSIQERRPQLPQVLQHVRLPLLSPKFL<br>VGTVGSDPLIKSDEECRDLVDEAKNYLLLPQERPLMQGPRTRPRKPIRCGE |
| March5$_{1-278}$ | MPDQALQQMLDRSCWVCFATDEDDRTAEWVRPCRCRGSTKWVHQACLQR<br>WVDEKQRGNSTARVACPQCNAEYLIVFPKLGPVVYVLDLADRLISKACPFAAA<br>GIMVGSIYWTAVTYGAVTVMQVVGHKEGLDVMERADPLFLLIGLPTIPVMLILG<br>KMIRWEDYVLRLWRKYSNKLQILNSIFPGIGCPVPRIPAEANPLADHVSATRILC<br>GALVFPTIATIVGKLMFSSVNSNLQRTILGGIAFVAIKGAFKVYFKQQQYLRQAH<br>RKILNYPEQEEA |
| SPOP$^{mut}$ [4] | SVNISGQNTMNMVKVPECRLADELGGLWENSRFTDCCLCVAGQEFQAHKAIL<br>AARSPVFSAMFEHEMEESKKNRVEINDVEPEVFKEMMCFIYTGKAPNLDKMA<br>DDLLAAADKYALERLKVMCEDALCSNLSVENAAEILILADLHSADQLKTQAVDF<br>INYHASDVLETSGWKSMVVSHPHLVAEAERSLASAQCPFLGPPRKRLKQ |
| **Controls** | |
| mCherry<br><br>(3xHA-T2A-mCherry) | YPYDVPDYAYPYDVPDYAYPYDVPDYAgssggsgEGRGSLLTCGDVEENPG↑P<br>ggsgVSKGEEDNMAIIKEFMRFKVHMEGSVNGHEFEIEGEGEGRPYEGTQTAK<br>LKVTKGGPLPFAWDILSPQFMYGSKAYVKHPADIPDYLKLSFPEGFKWERVM<br>NFEDGGVVTVTQDSSLQDGEFIYKVKLRGTNFPSDGPVMQKKTMGWEASSE<br>RMYPEDGALKGEIKQRLKLKDGGHYDAEVKTTYKAKKPVQLPGAYNVNIKLDI<br>TSHNEDYTIVEQYERAEGRHSTGGMDELYK* |
| αGFP no ligase<br><br>(3xHA-**αGFP**-T2A-mCherry) | YPYDVPDYAYPYDVPDYAYPYDVPDYAgsdqVQLVESGGALVQPGGSLRLSC<br>AASGFPVNRYSMRWYRQAPGKEREWVAGMSSAGDRSSYEDSVKGRFTISR<br>DDARNTVYLQMNSLKPEDTAVYYCNVNVGFEYWGQGTQVTVSSgssgsgEGR<br>GSLLTCGDVEENPG↑PggsgVSKGEEDNMAIIKEFMRFKVHMEGSVNGHEFEIE<br>GEGEGRPYEGTQTAKLKVTKGGPLPFAWDILSPQFMYGSKAYVKHPADIPDY<br>LKLSFPEGFKWERVMNFEDGGVVTVTQDSSLQDGEFIYKVKLRGTNFPSDGP<br>VMQKKTMGWEASSERMYPEDGALKGEIKQRLKLKDGGHYDAEVKTTYKAKK<br>PVQLPGAYNVNIKLDITSHNEDYTIVEQYERAEGRHSTGGMDELYK |
| SPOP E3 ligase only<br><br>(3xHA-**E3 ligase**-T2A-mCherry) | YPYDVPDYAYPYDVPDYAYPYDVPDYAgsdqSVNISGQNTMNMVKVPECRLA<br>DELGGLWENSRFTDCCLCVAGQEFQAHKAILAARSPVFSAMFEHEMEESKKN<br>RVEINDVEPEVFKEMMCFIYTGKAPNLDKMADDLLAAADKYALERLKVMCEDA<br>LCSNLSVENAAEILILADLHSADQLKTQAVDFINYHASDVLETSGWKSMVVSHP<br>HLVAEAYRSLASAQCPFLGPPRKRLKQSgssggsgEGRGSLLTCGDVEENPG↑P<br>ggsgVSKGEEDNMAIIKEFMRFKVHMEGSVNGHEFEIEGEGEGRPYEGTQTAK<br>LKVTKGGPLPFAWDILSPQFMYGSKAYVKHPADIPDYLKLSFPEGFKWERVM<br>NFEDGGVVTVTQDSSLQDGEFIYKVKLRGTNFPSDGPVMQKKTMGWEASSE<br>RMYPEDGALKGEIKQRLKLKDGGHYDAEVKTTYKAKKPVQLPGAYNVNIKLDI<br>TSHNEDYTIVEQYERAEGRHSTGGMDELYK* |

### Degradation motifs (DM)

| | |
|---|---|
| SPOP [5, 6] 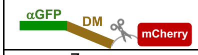  | DEVTSTTSSSTgsDEVTSTTSSSTgsDEVTSTTSS |
| VHL[7] 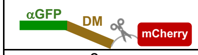  | ALAPYIPgssgsgALAPYIPgssgsgALAPYIP |
| CHIP [8] 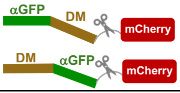  | LWWPDggggsgsLWWPDggggsgsLWWPDgsgsGSGPTIEEVDgsgsGSGPTIEEVDgsgsGSGPTIEEVD |
| βTrCP [9] 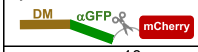  | AEGDTEEDDGFVDIgsAEGDTEEDDGFVDIgsAEGDTEEDDGFVDI |
| KLHL20 [10] 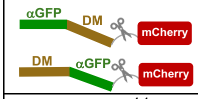  | LLAMNLGLPDLVAKYNTSNGAgsgsLLAMNLGLPDLVAKYNTSNGAgsgsLLAMNLGLPDLVAKYNTSNGA |
| KLHDC2 [11] 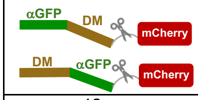  | HLRGSPPPMAGGgsgsHLRGSPPPMAGGgsgsHLRGSPPPMAGG |
| UBR [12] 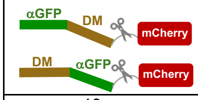  | KIGLGRQKPPKATKggggsgsKIGLGRQKPPKATKgsgsgsKIGLGRQKPPKATK |
| MC2 [13] 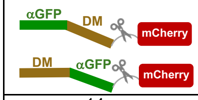  | SDKPLHRYVGFQCGggggsgsSDKPLHRYVGFQCGgsgsgsSDKPLHRYVGFQCG |
| ODC [14] 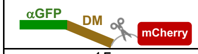  | SHGFPPEVEEQDDGTLPMSCAQESGMDRHPAACASARINV |
| UBL1 [15] 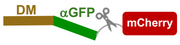  | MQVTLKTLQQQTFKIDIDPEETVKALKEKIESEKGKDAFPVAGQKLIYAGKILNDDTALKEYKIDEKNFVVVMVTKPKAVSTP |
| UBL2 [15] 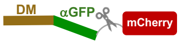  | MKSTFKSEYPFEKRKAESERIADRFKNRIPVICEKAEKSDIPEIDKRKYLVPADLTVGQFVYVIRKRIMLPPEKAIFIFVNDTLPPTAALMSAIYQEHKDKDGFLYVTYSGENTFGRggggsggggsMKSTFKSEYPFEKRKAESERIADRFKNRIPVICEKAEKSDIPEIDKRKYLVPADLTVGQFVYVIRKRIMLPPEKAIFIFVNDTLPPTAALMSAIYQEHKDKDGFLYVTYSGENTFGRggggsggggsMKSTFKSEYPFEKRKAESERIADRFKNRIPVICEKAEKSDIPEIDKRKYLVPADLTVGQFVYVIRKRIMLPPEKAIFIFVNDTLPPTAALMSAIYQEHKDKDGFLYVTYSGENTFGR |
| $(UBL)_3$ [15, 16] 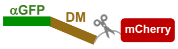  | MDLTISNELTGEIYGPIEVSEDMALTDLIALLQADCGFDKTKHDLYYNMDILDSNRTQSLKELGLKTDDLLLIRGKISNSgssgsgMSLNIHIKSGQDKWEVNVAPESTVLQFKEAINKANGIPVANQRLIYSGKILKDDQTVESYHIQDGHSVHLVKSQPKgssgsgMQVTLKTLQQQTFKIDIDPEETVKALKEKIESEKGKDAFPVAGQKLIYAGKILNDDTALKEYKIDEKNFVVVMVTKPKAVSTP |
| LIR1 [17] 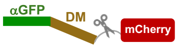  | FYVEESDNCSGGDDDWTHLSSKEVDPggggsgsESDNCSGGDDDWTHLSSKEVDPgsgsESDNCSGGDDDWTHLSSKEVDP |
| LIR2[18] 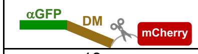  | VEEEWVIVSDEEIEEARQKAPLEITEYVEEEWVIVSDEEIEEARQKAPLEIT EY |
| LIR3 [18] 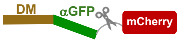  | VEEEWVIVSDEEIEEARQKAgsgsVEEEWVIVSDEEIEEARQKA |

| | |
|---|---|
| **Target-binding motifs (TBM)** | |
| $Ad_{BCL-xL}$ [19] | SENSLEIEELARFAVDEHNKKENALLEFVRVVKAKEQMFSWLDWEETMYYLTLEAKDGGKKKLYEAKVWVKPALLWSPHGNFKELQEFKPV |
| BAD [20] | NLWAAQRYGRELRRMSDEFVDSFKK |
| BAD-HRK [21] | SAAQLYAARLKAFGDEFHQ |
| HRK [21] | SAAQLTAARLKALGDELHQRTMW |
| $Ad_{BCL-xL}^{mut}$ W41A | SLEIEELARFAVDEHNKKENALLEFVRVVKAKEQMFSWLDAEETMYYLTLEAKDGGKKKLYEAKVWVKPALLWSPHGNFKELQEFKPV |
| $BAD^{mut}$ | GARLAQLKQERAALEQRLAALDQEIAALEWQIQSDPRKKQLEQKLLELQAERLRLIGDIVNLDQQILNLEAG |
| **Multimerization domains** fused at the N-terminus to a FLAG tag and BAD peptide (for BCL-$x_L$ binding); varying location of degradation motif (DM) indicated in Figure 6. | |
| FLAG-**BAD** | DYKDDDDKNLWAAQRYGRELRRMSDEFVDSFKKgsgss |
| apCC-Di [22] 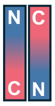  | G QLEQELAALDQQIAALKQRRAALKWQIQG |
| CC-Tri 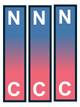  | GEIAAIKQEIAAIKKEIAAIKWEIAAIKQG |
| CC-Tet 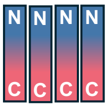  | GELAAIKQELAAIKKELAAIKWELAAIKQG |
| apCC-Tet 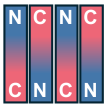  | GQLEEIAQQLEEIAKQLKKIAWQLKKIAQG |
| **Additional controls** | |
| FLAG-**BAD**-CC-Tet-T2A-mCherry | DYKDDDDKNLWAAQRYGRELRRMSDEFVDSFKKgsgssGELAAIKQELAAIKKELAAIKWELAAIKQGgssggsEGRGSLLTCGDVEENPG↑PggsgVSKGEEDNMAIIKEFMRFKVHMEGSVNGHEFEIEGEGEGRPYEGTQTAKLKVTKGGPLPFAWDILSPQFMYGSKAYVKHPADIPDYLKLSFPEGFKWERVMNFEDGGVVTVTQDSSLQDGEFIYKVKLRGTNFPSDGPVMQKKTMGWEASSERMYPEDGALKGEIKQRLKLKDGGHYDAEVKTTYKAKKPVQLPGAYNVNIKLDITSHNEDYTIVEQYERAEGRHSTGGMDELYK* |
| FLAG-CC-Tet-ODC-T2A-mCherry | DYKDDDDKgsgssGELAAIKQELAAIKKELAAIKWELAAIKQGggggsSHGFPPEVEEQDDGTLPMSCAQESGMDRHPAACASARINVgssggsEGRGSLLTCGDVEENPG↑PggsgVSKGEEDNMAIIKEFMRFKVHMEGSVNGHEFEIEGEGEGRPYEGTQTAKLKVTKGGPLPFAWDILSPQFMYGSKAYVKHPADIPDYLKLSFPEGFKWERVMNFEDGGVVTVTQDSSLQDGEFIYKVKLRGTNFPSDGPVMQKKTMGWEASSERMYPEDGALKGEIKQRLKLKDGGHYDAEVKTTYKAKKPVQLPGAYNVNIKLDITSHNEDYTIVEQYERAEGRHSTGGMDELYK* |

**Table S2. Natural and synthetic binding partners for BCL-x$_L$ and αGFP nanobody**

| Name | $K_d$ (nM) | $IC_{50}$ (nM) | Reference |
|---|---|---|---|
| vhhGFP4 | 1.4 | | [23] |
| Ad$_{BCL\text{-}xL}$ | 39 ± 1 (ITC) | 393 ± 54 (FA) | [19] |
| BAD$_{109\text{-}127}$ | 119 ± 23 (FA) | 200 ± 10 (FA) | [20] |
| HRK$_{28\text{-}50}$ | 38 ± 6 (FA) | 2200 ± 100 (FA) | [21] |
| BAD-HRK | | 900 ± 100 (FA) | [21] |

Technique used is indicated in parentheses (either isothermal titration calorimetry (ITC) or fluorescence anisotropy (FA)).

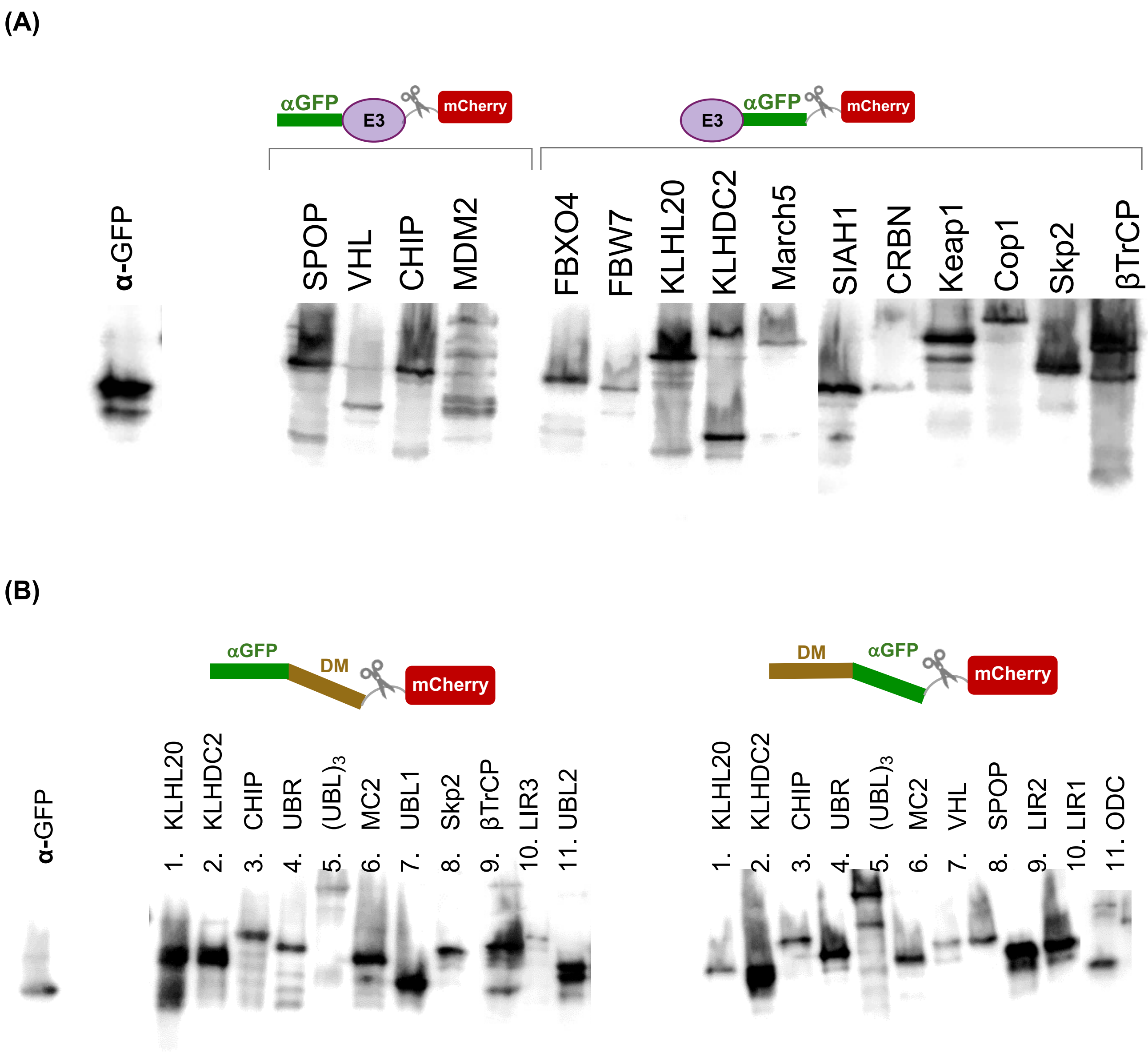


**Figure S2. Western blot of HA-tagged degraders probed by anti-HA high affinity monoclonal antibody (#45-11867423001, Roche). (A)** αGFP-based chimeric E3s and **(B)** αGFP-degron fusions.

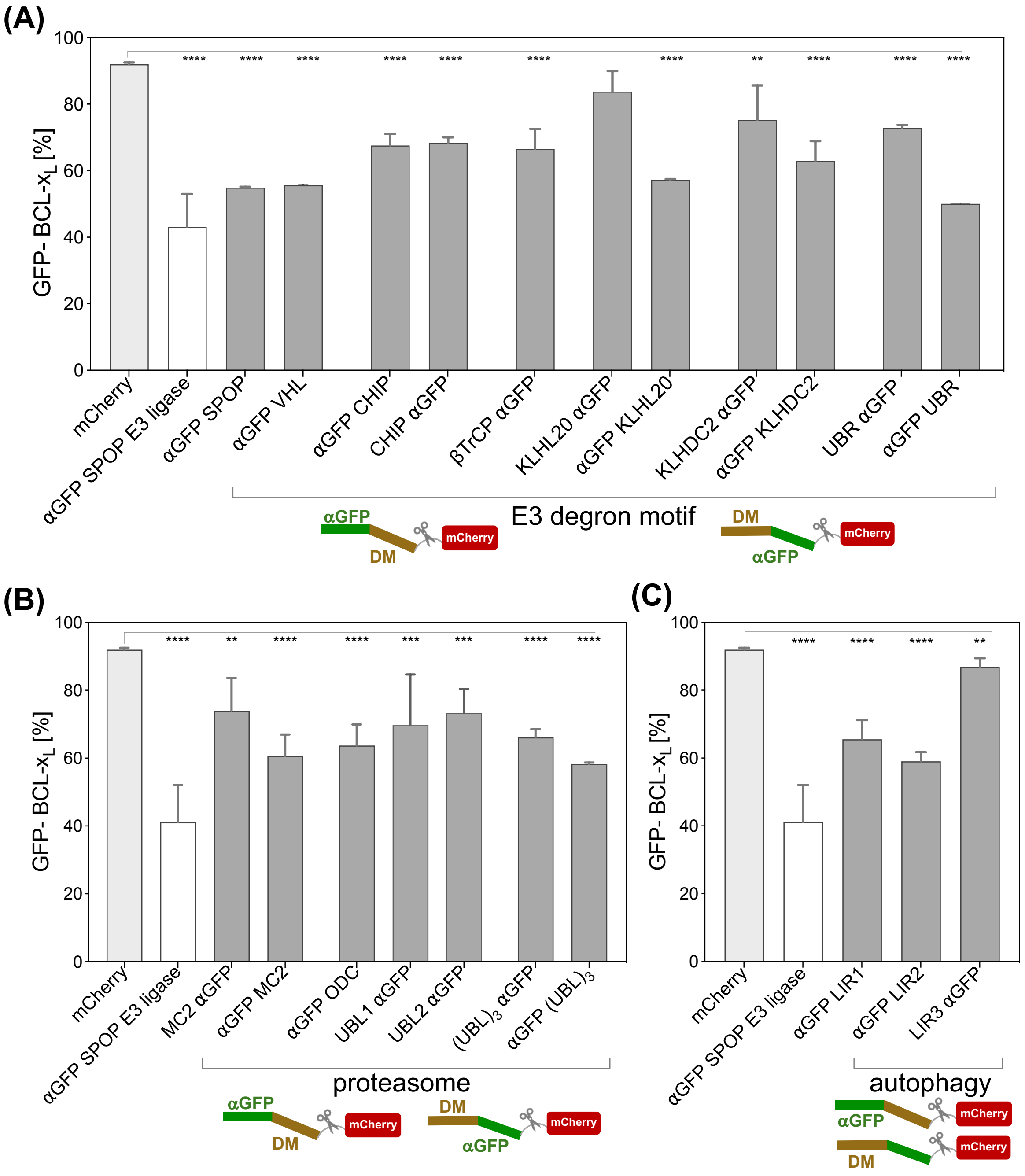


**Figure S3. GFP-BCL-x$_L$ is degraded by αGFP-degron fusions.** Flow cytometry analysis of HEK293T cells constitutively expressing GFP-BCL-x$_L$ 48 hours post transfection with **(A)** degraders composed of degrons for E3s, **(B)** degrons targeting directly to the proteasome, and **(C)** degrons targeting to the autophagy-lysosome pathway.

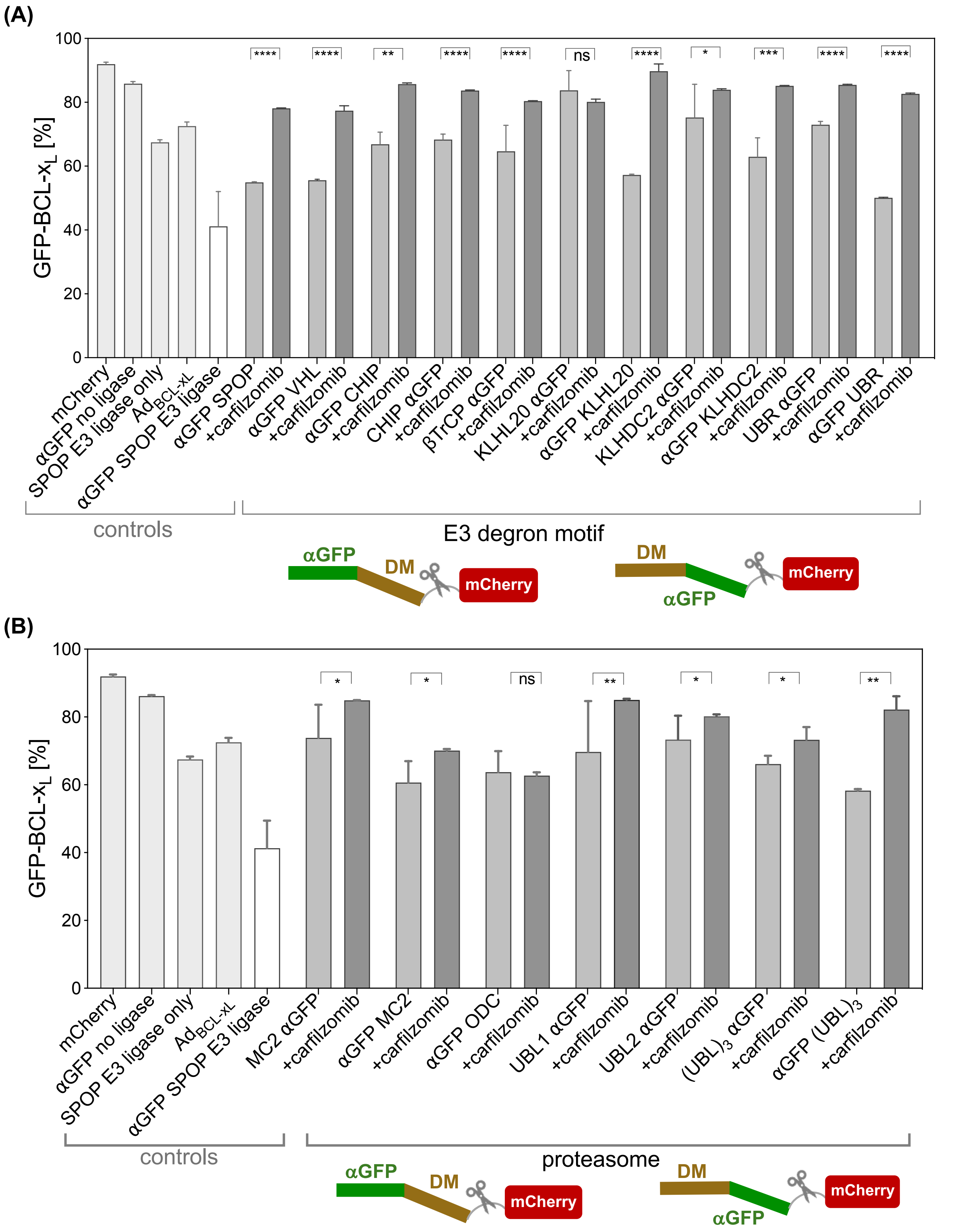


**Figure S4. Flow cytometry analysis of HEK293T cell line constitutively expressing BCL-x$_L$ 48 hours post transfection with αGFP-degron fusions upon treatment with carfilzomib.** Cells were treated with 20 μM carfilzomib for 24 hours prior to measurement (Carfilzomib (PR-171), #A1933, APExBIO). (**A**) Degrons for E3s; **(B)** degrons for targeting to the proteasome. Statistics were calculated using Welsch's t test.

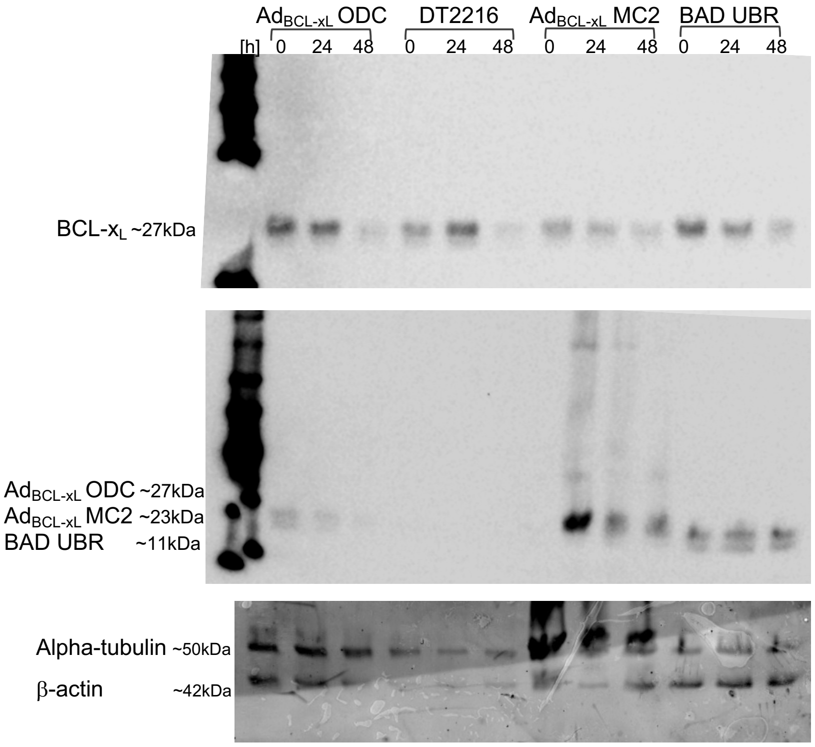


**Figure S5. Western blot showing representative examples of BCL-x$_L$ and degrader levels 24 hours and 48 hours post degrader transfection.** anti-BCL-x$_L$ (54H6) Rabbit mAb (HRP Conjugate) (#24780, Cell Signaling) was used for BCL-x$_L$ detection, anti-HA high affinity monoclonal antibody (#45-11867423001, Roche) for detection of: Ad$_{BCL-xL}$ ODC, Ad$_{BCL-xL}$ MC2 and BAB UBR, and Alpha Tubulin Monoclonal antibody (# 66031-1-Ig, Proteintech) and Actin Antibody (I-19)(sc-1616, Santa Cruz) for loading controls, respectively.

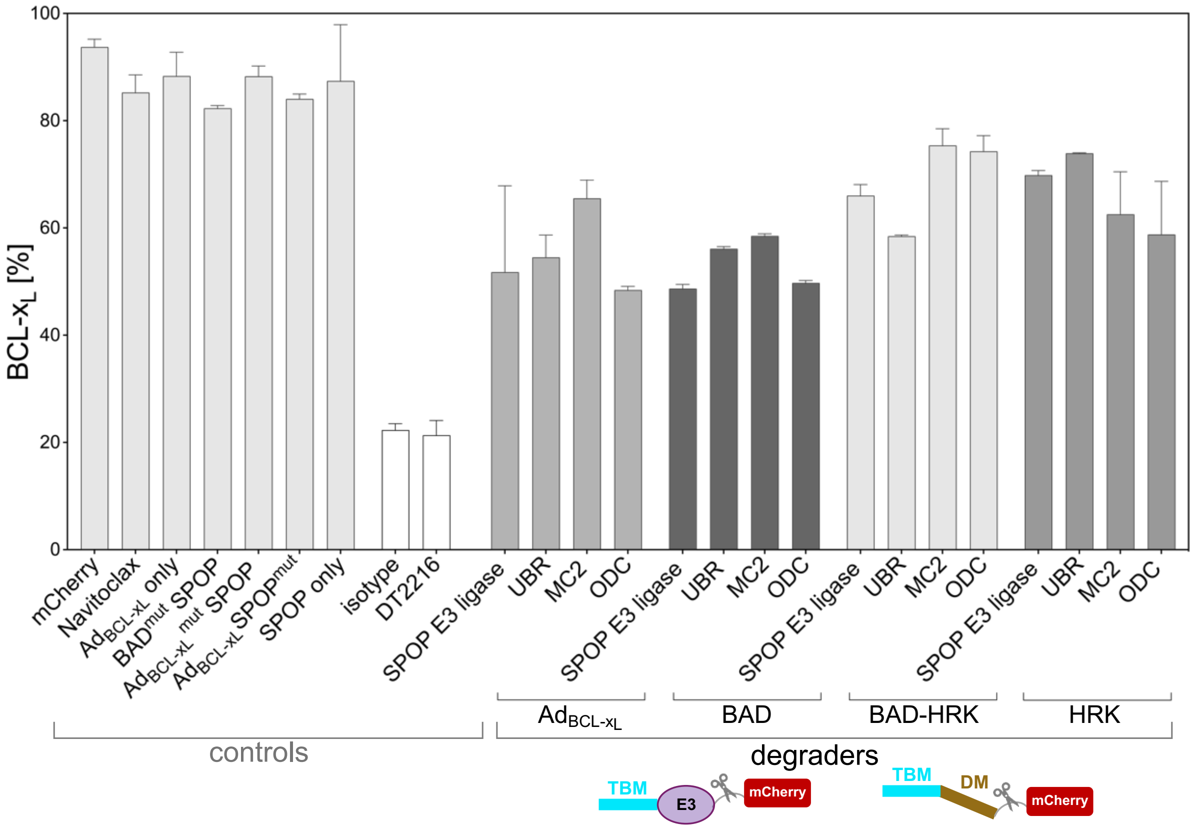


**Figure S6. Additional control molecules in endogenous BCL-x$_L$ degradation assay.** Navitoclax and DT2216 are shown as controls. Cells were treated with Navitoclax at 3.3 $\mu$M or DT2216 at 1 $\mu$M for 48 hours before flow-cytometry measurements (#HY-10087, MedChemExpress and #37311, Cayman Chemical, respectively). A Rabbit IgG Isotype control, Alexa Fluor 488 Conjugated, was included. Sequences of controls and degraders are in Table S1. Results are from at least 3 experiments and depicted as the mean ± SD.

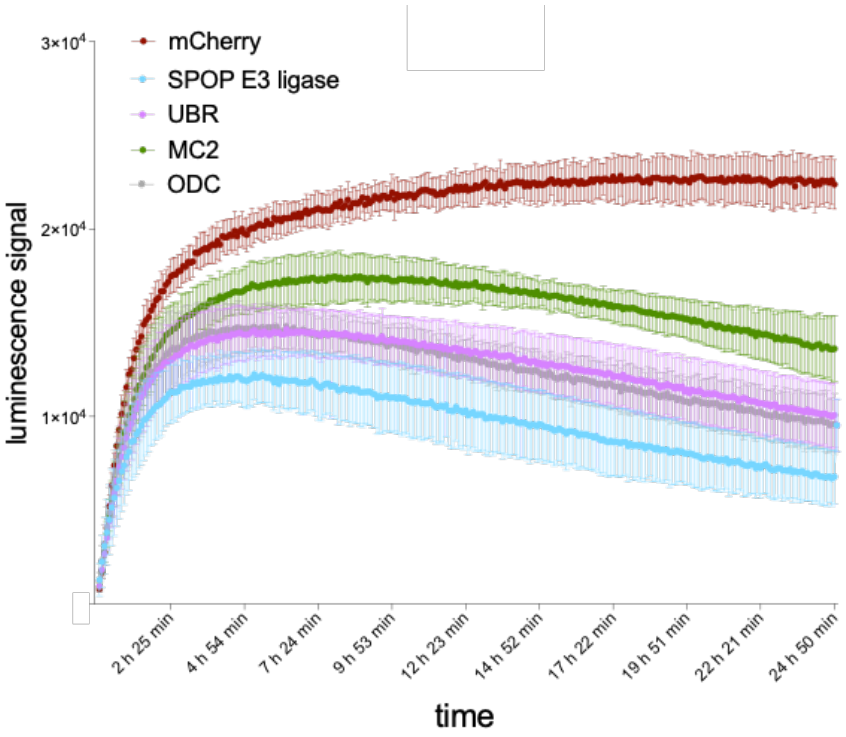


**Figure S7. Representative examples of time dependence of degradation of Nanoluc- BCL-x$_L$ monitored by luminescence.** Cells were transfected with degraders and luciferase substrate (endurazine) was added at the same time (corresponding to t0). Degraders shown are those containing BAD as the BCL-x$_L$ binder. SPOP indicates chimeric E3, all others are BAD-degron fusions.

**(A)**

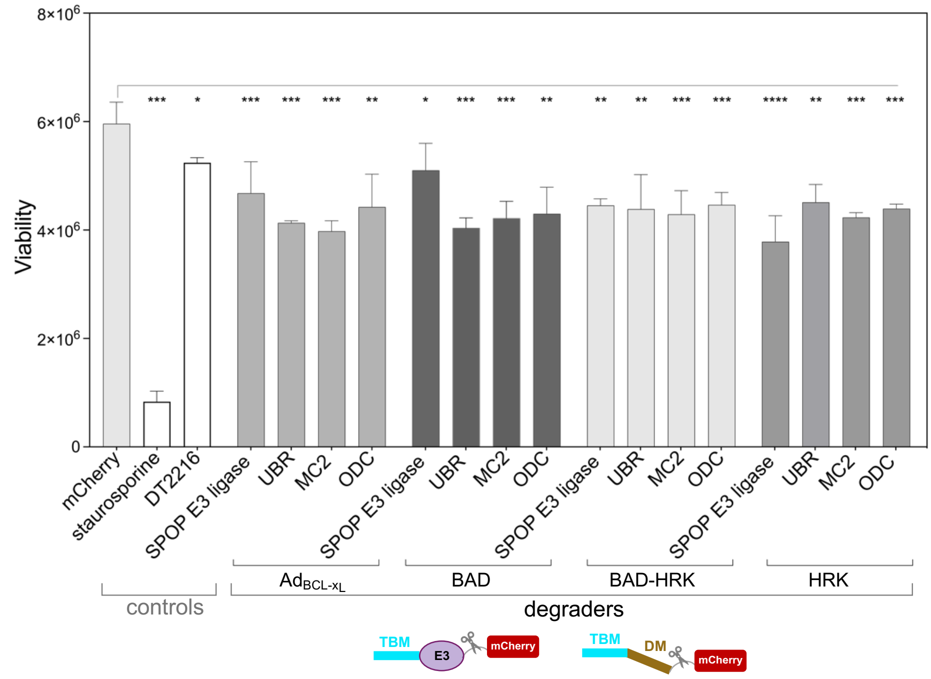


**(B)**

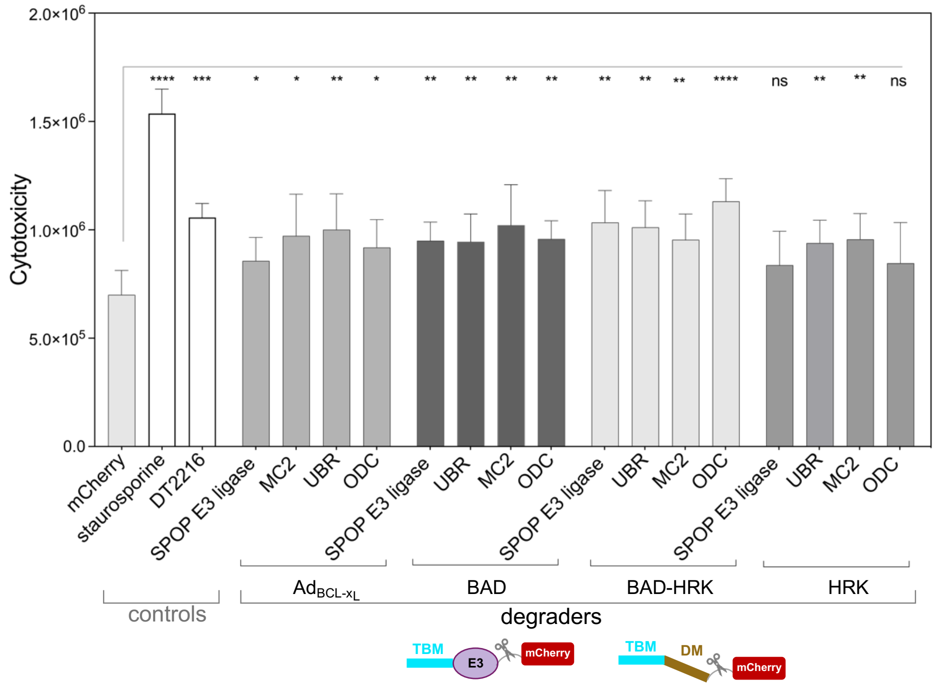


**Figure S8. Cell viability (A) and cytotoxicity assays (B).** Staurosporine as an apoptosis inducer was included as a control; cells were subjected to 24-hours' treatment with 10 nM staurosporine prior to measurement [24].

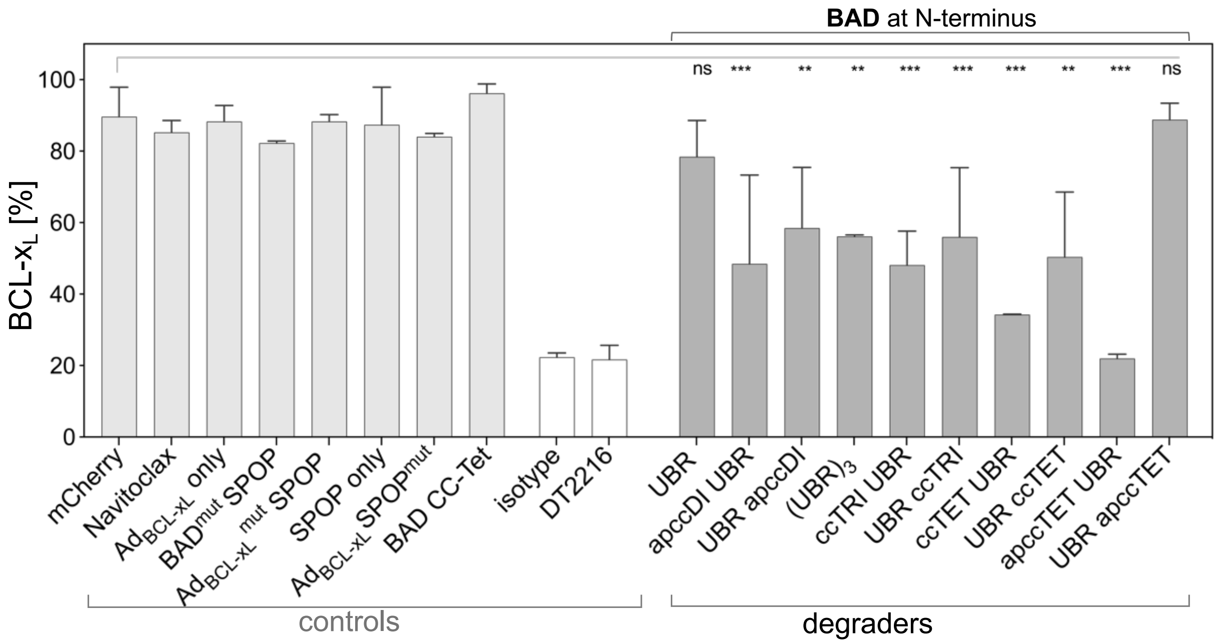


**Figure S9. Degradation of endogenous BCL-x$_L$ using multivalent degraders.** Degradation was measured by flow cytometry 48 hours post transfection with the degraders.

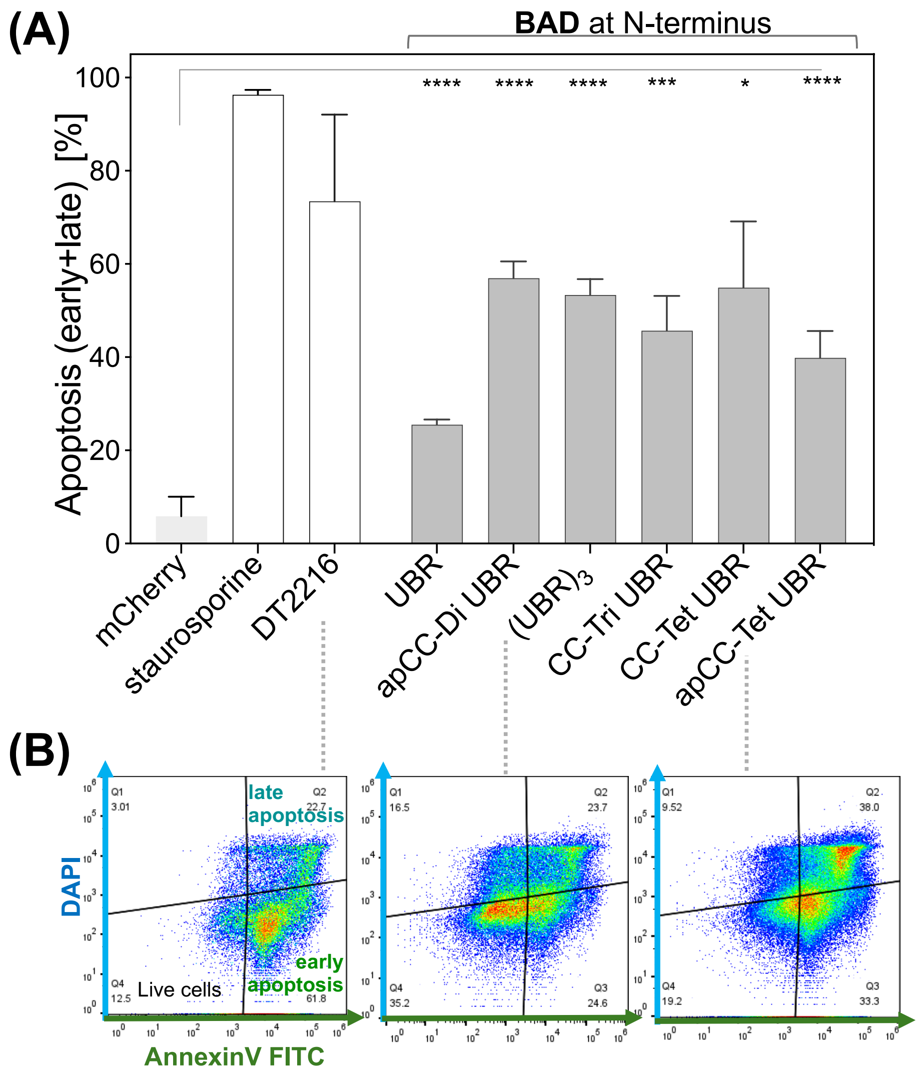


**Figure S10. Induction of apoptosis using multivalent degraders. (A**) Induction of apoptosis 48 hours post transfection of A549 cells with degraders, PROTAC (DT2216), or staurosporine control; (**B**) representative flow cytometry of FITC-labelled Annexin V and DAPI staining (BioLegend and BD Pharmingen, respectively).
